# Linguistic structure and probability are jointly encoded in high gamma power

**DOI:** 10.64898/2026.07.21.739808

**Authors:** Sophie Slaats, Alexis Hervais-Adelman

**Affiliations:** Department of Basic Neurosciences, Faculty of Medicine, University of Geneva, Geneva, Switzerland

**Keywords:** high gamma power, electrocorticography, language comprehension, syntax, surprisal, speech, multivariate temporal response functions

## Abstract

During speech comprehension, the brain dynamically infers a hierarchy of increasingly abstract representations from the sensory input. An important step in the inferential hierarchy is the combination of words to form phrases and sentences. Whether this process is driven primarily by statistical patterns in the linguistic input, or by a mechanism that combines words into hierarchical representations, is a subject of considerable debate that has regained importance with the arrival of large language models. This study investigates whether local cortical activity (high gamma power; 70-150 Hz) from intracranial recordings is jointly modulated by lexical probability and syntactic structure; and whether lexical probability affects the inference of syntactic structure. To this end, an open dataset of electrocorticography recordings is analyzed with multivariate temporal response functions and a model comparison approach. The results indicate that high gamma power is sensitive to multi-word estimates of constituency structure and lexical probability estimates, both in isolation and jointly. The temporal response functions suggest that syntactic structure building depends on interregional communication between regions connected through dorsal- and ventral streams. Furthermore, the study provides evidence that bottom-up syntactic information is less likely to be encoded by neural populations that strongly code for lexical probability measures, while top-down syntactic information shares neural resources with lexical uncertainty. We suggest that lexical uncertainty modulates the weighting of anticipatory structural information. With this, the current study supports models that suggest that cues are leveraged flexibly in a feed-forward and feed-back fashion during speech comprehension.

## 1. Introduction

During language comprehension, the brain infers a hierarchy of increasingly abstract representations from the sensory input, from phonemes to the meaning of the message. This is viewed as a dynamical process in which representations can cue each other both in a feed-forward and feed-back manner (1,2). An important step in the inferential hierarchy is the combination of words to create interpretable phrases and sentences. High frequency neural activity in the gamma band (70-150 Hz) has been suggested to play a critical role in this process (2–9). Whether it is driven primarily by the distributional statistics of tokens in linguistic input, or by a mechanism that combines words into abstract hierarchical representations is a subject of considerable debate (10–15). In this study, therefore, we explore whether these sources of information are jointly represented within high gamma power (70-150 Hz) during spoken language comprehension.

### 1.1. Lexical probability and syntactic structure in neural dynamics

There is much evidence for the separate and joint use and modulation of neural activity by statistical and structural information (for example, 15–20). In a seminal study, Ding and colleagues (21) showed that neural activity reflects linguistic structure by displaying a power-increase that the rate of occurrence of that unit. In that study, results revealed a 1 Hz power increase as a response to the occurrence of four-word sentences and a 2 Hz increase for two-word phrases – with the (monosyllabic) words presented at a 4 Hz rate. Subsequently, Batterink and Paller (22) showed that similar signatures of representations can be elicited on the basis of transitional probability alone. In a statistical learning paradigm, the authors presented participants with two conditions of syllables; a random condition, and a condition in which syllables could form trisyllabic ‘non-words’ based on the transitional probability between syllables. The results revealed that power at 1.1 Hz – the rate of occurrence of the non-words – increased as the participants learned the transitional probabilities in the input.

These sets of findings together indicate that the brain creates latent representations using both statistical patterns in the input (probability) and knowledge of the structure of language (syntax). The studies show that statistical patterns and linguistic information can exist and be leveraged in isolation. In naturalistic contexts, however, linguistic structure and statistical patterns (probability) are related: the presence of linguistic structure drives specific statistical regularities in language (14,23), and these statistical regularities can help the perceiver to find structure in the input (24–28). Indeed, MEG studies have shown that slow neural dynamics jointly encode structural and statistical properties (15,29) and that these two types of information can interact: lexical probability affects the time-course of structure building processes captured by delta-band activity, suggesting that probability plays a role in structure building (30); and the representation of lexical probability in the neural signal is affected by the syntactic status of a word (31). This set of findings indicates that probabilistic and syntactic information may be exploited dynamically during speech comprehension (14). Besides low-frequency activity, the high gamma band (70-150 Hz) has been proposed to play an important role in speech processing (32) and structure building in particular (2–9). Activity in this frequency band is commonly assumed to reflect the average firing rates and synchrony of the local neural population (33–35). Concretely, high gamma power (HGP) has been suggested to reflect the active process of binding (6), the semantic composition of language-specific concepts (3,9), and the application of grammatical rules or stored lexical knowledge to infer a higher-level structure (2,36). Any of these processes may depend on or elicit modulations of patterns of inhibition, a process to which gamma activity broadly has been related (2,37,38).

Indeed, HGP can reflect syntactic structure. Ding and colleagues (21) also analysed power in the high gamma band (70-200 Hz) recorded with electrocorticography (ECoG). This analysis revealed that HGP responded similarly to the sentences, phrases, and word lists as low-frequency activity, namely: by an increase in the amplitude of the HGP signal at the frequency corresponding to the rate of occurrence of the linguistic unit in electrodes in the middle and posterior superior temporal gyrus (STG) and left inferior frontal gyrus (IFG). And similarly to the effects shown with M/EEG, subsequent studies revealed that power increases in HGP at the corresponding rate of occurrence can be elicited using a classical, syllable-based statistical learning paradigm, too (39,40). This suggests that HGP plays a role (either causal or consequential) in both syntactic structure building and in segmentation using transitional probability information.

### 1.2. Syntactic structure in high gamma power: Multi-unit structure or semantic composition?

Following Ding et al. (21), there have been several studies showing that HGP increases as structure accrues, both in reading (41–44) and in speech comprehension (45–47), though whether this effect is driven by syntactic structure or semantic composition is debated (9). These studies broadly employed two types of experimental designs to isolate effects of structure building: condition-based and regression-based approaches. In the condition-based approaches, a frequently employed paradigm hinges on the comparison of stimuli that (do not) contain specific linguistic aspects of the stimulus, e.g. comparisons among full sentences (contain structure and meaning); jabberwocky sentences (‘sentences’ of pseudowords; contain structure but no meaning); word lists (contain meaning, but no structure); and lists of pseudowords (contain neither). Regression-based approaches use parsing models to specify how structure is constructed when incoming words are added, creating ‘nodes’ in a tree structure graph of the sentence (see Figure 2 below for an example). The structures described in this paper are *constituency structures:* tree structures that describe syntactic relations hierarchically based on phrase structure rules. Syntactic *dependencies* may also be used. Linking functions are discussed in more detail elsewhere (e.g. in 18,48).

Fedorenko and colleagues (42) showed that cortical HGP (70-170 Hz) recorded with ECoG increases monotonically over the course of reading a sentence, an effect that diminished as linguistic information was removed (in order: word lists, jabberwocky sentences, pseudoword lists). The regression-based approach used by Nelson and colleagues (43) revealed that the average HGP (70-150 Hz) between 200 to 500 milliseconds post word onset increases with the number of open nodes in a structure, and drops when nodes are closed. Using the same conditions as Fedorenko et al. (42), Woolnough and colleagues (44) reported the increase in HGP as the sentence progresses, but failed to find the effects of open nodes and node closures reported by Nelson and colleagues (43). The study suggested that two spatiotemporally distinct networks are involved in structure building during reading: a network of IFG and the middle temporal gyrus (MTG), which shows the increase as structure accrues; and a network that engages IFG and STG and shows greater activation for words in word lists than for words in sentences.

Murphy and colleagues (46) compared adjective-noun, pseudoword-noun, and adjective-pseudoword combinations. This revealed that sensors in posterior superior temporal areas and IFG (pars triangularis) showed responsiveness either to ‘lexicality’ (pseudoword vs. word), with higher gamma band activity (70-150 Hz) for pseudowords; or to ‘compositionality’ (adjective-noun vs. the combined pseudoword trials), with higher gamma activity for the compositional phrases than the pseudoword combinations. The degree of overlap was very small, with in total only three electrodes displaying effects for both contrasts. In a different study, Murphy et al. (47) then compared combinations that manipulated subject-verb agreement: pseudoword-verb (‘trab run’), pronoun-verb *grammatical* (‘they run’) and pronoun-verb *ungrammatical* (‘they runs’; agreement error). The results showed effects of composition (pseudowords vs. grammatical), HGP was increased for compositional phrases; and of agreement error (grammatical vs. ungrammatical), HGP was higher for ungrammatical phrases than for ungrammatical phrases. Both effects were found in electrodes in posterior temporal areas and IFG, but as before, only a small number of closely neighboring sites were sensitive to both contrasts. The authors suggest that these results potentially support a framework of initial, rapid structure-generation in posterior superior temporal sulcus (STS) and posterior MTG, which is followed by a more semantically demanding interpretation process in IFG. Taken together, these two studies suggest that populations of neurons whose activity is captured by one electrode are relatively selective: most contacts are sensitive to either lexicality or grammatical contrasts, and not both.

### 1.3. Surprisal and entropy in high gamma power

Besides linguistic structure, Nelson et al. (43) also considered effects of trigram lexical probability measures on HGP, namely surprisal and entropy. Surprisal is the (log) probability of a word given the context (two words in the case of a trigram model). If surprisal is high, the word was unexpected (either unpredictable or unpredicted). Entropy is the weighted average of all surprisal values of the words that can appear in the target position and captures uncertainty. High entropy means high uncertainty. Nelson and colleagues showed that entropy was inversely related to HGP in superior and middle temporal areas; in other words, HGP decreased for increased uncertainty. In the same areas, there was a positive effect of surprisal, which means that HGP increased as word probability decreases. Adding surprisal and entropy as cofactors did not change the effects for the syntactic predictors, providing the first evidence that these factors are jointly modulating local, high frequency neural activity.

There have since been analyses of linguistic encoding in high gamma band activity using surprisal and entropy (and other measures) obtained from large language models (LLMs) as the features of interest (49–52). These estimates are frequently taken to reflect probabilistic processing in the human brain (53). By analyzing a dataset of naturalistic language comprehension, Goldstein and colleagues (50) showed that HGP (70-200 Hz) during language comprehension can be modeled by a combination of entropy and surprisal derived from LLMs such as GPT2. The correlations between the probability values and cortical activity were largest just before word onset for entropy, and at 400 milliseconds post word onset for surprisal. These effects were found in superior temporal and inferior frontal regions. Importantly, using the same dataset, Zou and colleagues (31) showed that probability and syntax are not encoded independently. By dividing words based on the position in a phrase – either phrase-initial or not –, the authors showed that the correlation between surprisal and high gamma activity interacted with the syntactic status of a word. Specifically, the correlation between GPT2-derived surprisal and high gamma activity was lower for phrase-initial words than for phrase-internal words. These effects existed in STG and IFG.

### 1.4. The present study

Taken together, these results suggest that HGP encodes both lexical probabilistic and structural information. HGP increases as structure accrues over the course of a sentence, and it can be modulated by lexical status (word/pseudoword) and probability (surprisal/entropy); whether the structural effects are the result of multi-word syntactic measures like node counts, or whether they reflect a process of semantic composition remains unknown. Across studies, the effects were found in several areas of the cortex commonly associated with language comprehension, notably the left temporal lobe, mainly the posterior STG, and the left IFG. None of the studies found a particular spatial location for syntactic or lexical (probabilistic) processing.

While results from lower-frequency activity increasingly suggest that lexical probabilistic and syntactic information are actively integrated during language comprehension (1,15,30,31,54), and HGP holds theoretical importance for syntactic structure building (2,7,9), little is known about how these two sources of information are encoded in HGP (31,43). This study, therefore, aims to provide insight into how these features are *jointly* encoded in cortical activity. We use the term ‘joint encoding’ to refer to the case where multiple features are encoded by the neural population whose activity is recorded by a single electrode (diameter of 5 mm^2^). Furthermore, this study aims to investigate if and how probabilistic information at the lexical level affects the neural encoding of syntactic structure, indicating interactions between representations at different levels in the inference hierarchy. The study broadly addresses the following questions: (1) Does HGP activity encode lexical probability and syntactic structure? And (2) does lexical probability affect HGP neural correlates of syntactic structure building? By using multivariate temporal response functions and cumulative model comparison in combination with naturalistic stimulation annotated for constituency structure, this study adds detail to previous findings in two domains: firstly, the analysis simultaneously estimates the contributions from multiple syntactic and probabilistic features; and secondly, the temporal response functions provide insight into the time-course of both feature types time-locked to word onset.

## 2. Materials and methods

Data was obtained from a published dataset of human intracranial EEG recordings collected at the University Medical Center Utrecht (UMC Utrecht, The Netherlands) (55). The full dataset contained data from 64 patients who underwent intracranial electrode implantation as part of surgical treatment for drug-resistant epilepsy. The data are available on OpenNeuro: https://openneuro.org/datasets/ds003688/. The code used for analysis can be found on Github: https://github.com/sslaats/ieeg-gamma-syntax. Other materials, including supplementary materials, are shared on the Open Science Framework: https://osf.io/vf2dg/.

### 2.1. Participants

In this study, we included a subset of 18 adults (18-52 years old, mean: 37 years 3 months, s.d.: 10 years 11 months; 12 female). These were selected according to the following criteria: the participants (1) were 18 years of age or older; (2) had a left-lateralized language network as per the clinical test conducted by the clinicians from the UMC Utrecht; (3) were implanted with left-lateralized ECoG grids; and (4) were right-handed. These criteria were used to ensure that the findings describe effects of language comprehension in a mature language system and are comparable to findings from the literature. The resulting selection had a total of 1377 electrodes (mean: 76 electrodes per patient, s.d.: 16), distributed across the left hemisphere in (pre)frontal, central, temporal and occipital regions as shown in in Figure 1.

**Figure 1.**
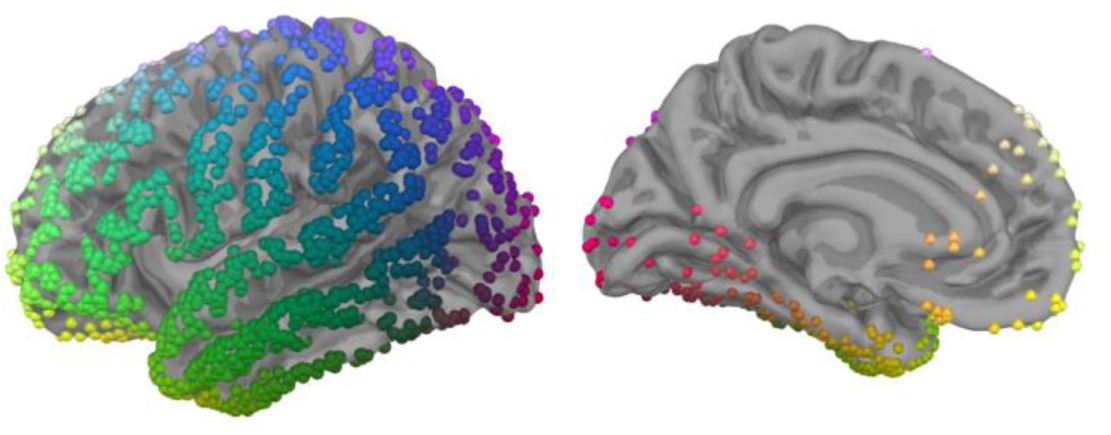
Electrode coverage of all analyzed electrodes (N=1377) projected onto fsaverage. RGB coloring reflects the 3D-positioning of the electrode.

### 2.2. Procedure

Participants were presented with the short film “Pippi on the run” (56), dubbed into the participants’ native language, Dutch. The film was shown in excerpts of 30 seconds, accompanied by either music (seven blocks) or speech (six blocks) from the movie. Each excerpt was unique. Originally, this task was designed for functional language mapping prior to surgical intervention. All participants provided informed consent to participate in the study and to publicly share their anonymized data (55). All procedures were approved by the Medical Ethical Committee of the UMC Utrecht in accordance with the Declaration of Helsinki (57).

### 2.3. iEEG and T1 preprocessing

Preprocessing was carried out using MNE-Python (58) version 1.8.0 in Python version 3.12.7. Channels marked as bad by the experimenters were dropped prior to processing. A notch filter was applied at 50 Hz and harmonics to remove line frequency interference, after which the data were re-referenced to the average. Since sampling rate varied across recordings, data were resampled to 512 Hz if the original sampling rate was higher. The data was epoched into segments of 30 seconds to extract the speech and music segments. Only the speech epochs were retained.

We analyzed broadband high gamma power (HGP), which was characterized using Morlet wavelets for 40 linearly spaced frequencies between 70 and 150 Hz (FFT-based convolution) and downsampled to 128 Hz using the *mne.time_frequency.tfr_morlet* function. The 40 resulting time-courses with a 128 Hz sampling rate were averaged to obtain broadband HGP. The time-courses were converted to decibel scale relative to the mean of the full recording (as in: 41).

To localize the electrode onto the individual participants’ native cortical surface for visualization and mapping into a common space (fsaverage), the cortical surface was reconstructed from the participants’ T1 structural MRI using Freesurfer’s (version 7.3.2) *recon_all* algorithm (59–61). The affine registration and symmetric diffeomorphic registration (SDR) between the participant’s and the fsaverage space was computed using the *mne.trasnforms.compute_volume_registration* function. The results were checked using the *mne.transforms.apply_volume_registration* function and visualization of the alignment. Before applying the affine and SDR transformations to the participants’ electrode locations, the locations provided with the dataset were transformed to the Freesurfer RAS space. To account for any changes in the participants’ neural surface since the acquisition of the T1 structural MRI scan, the coordinates were assigned the nearest vertex onto the participants’ individual pial surface using *scipy.spatial.cKDTree* (62 version 1.14.1). This step was skipped for two participants with lesions for whom this transformation led to noticeable distortions of the electrode locations (sub-06 and sub-24). The resulting electrode locations were then warped to the fsaverage cortical surface using the *mne.ieeg.warp_montage* function, which uses a montage created from the native electrode locations, the native MRI, the fsaverage MRI, the affine registration and the SDR. The resulting electrode locations were then assigned the nearest vertex onto the fsaverage pial surface, also using *scipy.spatial.cKDTree*.

### 2.4. Temporal Response Functions

To estimate how HGP relates to syntactic and probabilistic processes, we used (multivariate) temporal response functions (TRFs) to model the impact of a set of syntactic and lexical properties on the signal recorded at each electrode. In essence, TRFs are a multivariate linear regression approach. Using annotations of multiple aspects of the linguistic stimulus as features (see below), the TRF allows us to estimate how various aspects of language affect neural activity. The multivariate approach enables the estimation of effects for each feature separately. By lagging the features in time relative to the continuous neural signal, the TRF estimates modulation of these features at different points in time. The model returns a time-course of coefficients reflecting the encoding of a feature in the neural signal across time, in the present case, at the level of individual electrodes. In other words, an increase at a given time-lag for a feature reflects an increase in the associated brain response to this feature, at that electrode, and at the given time-lag after stimulus onset. Each feature has its own associated time-course of coefficients.

The equation (1) of the linear model is as follows:

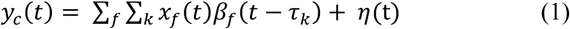

where {y_c_}_t_, {x_f_}_t_, {β_f_}_t_ represent the HGP signal of a given electrode c, the input feature f and its TRF coefficient at time lag τ_k_, respectively. {η}_t_ is Gaussian noise that accounts for variance not captured by the coefficients attributed to the features in the linear model. We used ridge regression to solve this equation, which means that the coefficients were estimated by minimizing the squared error between the actual HGP and the reconstructed HGP while keeping the norm of the coefficients low to avoid overfitting. The minimization problem is solved in closed form by the following equation:

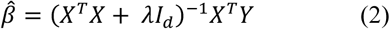

In this equation, Y denotes the measured signal in a matrix of real numbers of N samples by C channels; *β̂* is the set of beta coefficients (i.e., the estimated TRFs) organized in a matrix of real numbers with shape (K.F) x C, with K lags, F features for C channels; X is a matrix of real numbers containing all lagged feature time series with shape (K.F) x N, with K lags, F features of length N; λ is the regularization parameter and I_d_ is the identity matrix. The regularization parameter prevents overfitting (i.e., X^T^X not being full rank). In TRF models, this is often needed because linguistic features tend to be (auto)correlated, leading to numerically small eigenvalues.

We optimized regularization parameter λ using leave-one-out cross-validation over the six 30-second epochs, with 50 log-spaced values from 2^-2^ to 2^40^. We evaluated our model performance by computing the coefficient of determination R^2^ between the measured and reconstructed neural signal of the test epoch as the estimate of reconstruction accuracy. λ was chosen as follows. Firstly, we removed all electrodes for which the reconstruction accuracy never reached a value above 0. Secondly, we extracted the highest reconstruction accuracy values for each electrode per fold and computed the median across the remaining electrodes to obtain the value that worked best for most electrodes, resulting in six values (one per fold). We then chose the regularization parameter λ as the median of those six values. See Figure S1 (Supplementary Figures; https://osf.io/vf2dg/) for an example of the R^2^ values across folds for one participant and the chosen lambda value. The estimates of the beta coefficients and reconstruction accuracy that are reported and used for further analysis are the average beta coefficients and R^2^ values over all the folds. We chose this approach because of the short duration of the data. In this way, all epochs contribute to the beta coefficients and the reconstruction accuracy.

### 2.5. TRF feature design

To distinguish several sources of variance in the signal stemming from language comprehension, we constructed seven experimental features from the linguistic stimulus in three categories: base features, which were present in every model and are used to account for variance that could affect the results; and two sets of experimental features, namely lexical probability and syntactic features. Prior to the construction of the features, the provided transcription of the Dutch text was checked and corrected where necessary. This led to a slight reduction in the total number of words in the stimulus (N = 442 instead of N = 447). The word onset time-points provided with the dataset were adjusted accordingly.

#### 2.5.1. Base features

We constructed three base features: *speech envelope*, *word onset*, and *word frequency*.

The speech envelope feature was added to capture variance from early auditory processing to separate it from processing at the lexical and syntactic level. The speech envelope was computed by taking the absolute value of the Hilbert transform for each speech segment, and resampling this to 128 Hz to match the sampling rate of the (preprocessed) ECoG data.

The word onset feature was added to capture any response time-locked to word onset that was not accounted for by any of the other (more informative) features. To create the word onset feature, we used word onset time-points to create a time-course of zeros with values of one inserted at the sample that corresponded to the onset of a word. These time-series were convolved with a Gaussian kernel with a standard deviation of 15ms, for two reasons. Firstly, this accounts for any temporal inaccuracy of or uncertainty about the exact word onset time-points. Secondly, this process ensures that the feature requires a degree of regularization similar to the continuous speech envelope feature by increasing its autocorrelation.

The word frequency feature was used to capture decontextualized lexical processing. Like the word onset feature, this feature consisted of a time-series of zeros with values inserted at word onset. The values in this case were Zipf word frequency values obtained from the *wordfreq* package (63, version 3.1.1) with the *zipf_frequency* function for Dutch (see Van Heuven et al. (64) for the use of the Zipf metric). The resulting feature was equally convolved with a Gaussian kernel with a standard deviation of 15ms. Figure 2 shows an example of the resulting features.

**Figure 2.**
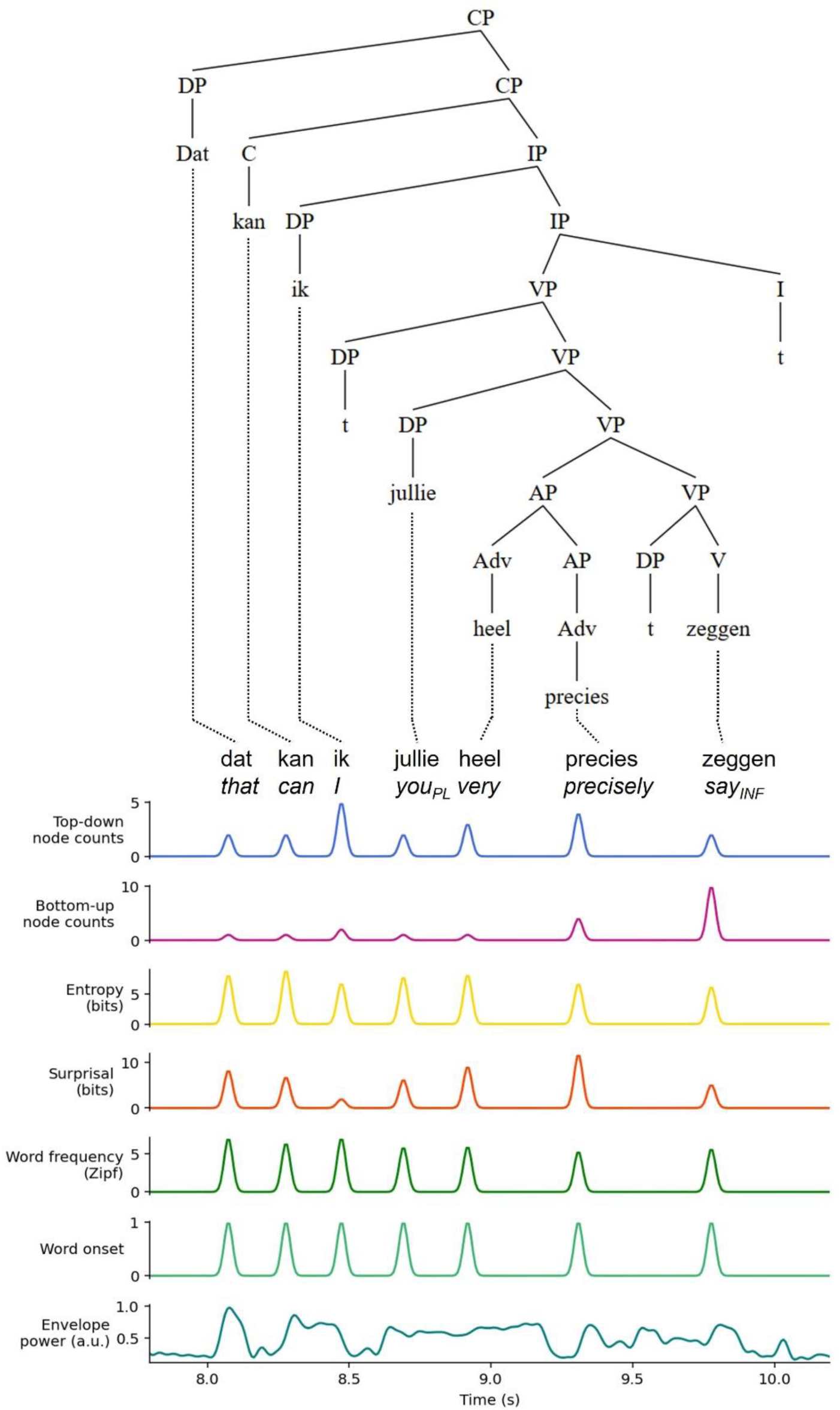
A parsed sentence from the stimulus and its corresponding features for the TRF models. The sentence reads, in Dutch: “dat kan ik jullie heel precies zeggen”, which translates to: “That, I can tell youPL very precisely.” Abbreviations: CP = complementizer phrase; DP = determiner phrase; IP = inflectional phrase; VP = verb phrase; AP = adverbial phrase; C = complementizer; I = inflection; Adv = adverb; V = verb.

#### 2.5.2. Lexical probability features

The lexical probability features were *surprisal* and *entropy.* Surprisal (*I*) quantifies the likelihood of a word occurring in context by taking the negative log probability *P* of the word (w_i_) given the preceding words (w_i-1_…w_i-n_); see the equation in (3) below.

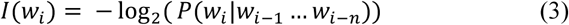

High values correspond to words that had a low likelihood or a high *surprise* given the context, and low values correspond to words that had a high likelihood given the context.

Entropy (*H*) quantifies uncertainty about the upcoming word based on the context. It is quantified by taking the weighted average of the surprisal values for all possible words (w_k_) that can occur in the position of word w_i_; see equation (4) below. When entropy values are high, the uncertainty about the next word is high. This happens when many words can occur in position *i*, and/or when all possible words have similar probability values. When one word is much more likely than others, entropy values decrease.

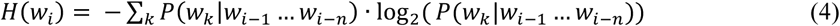

We obtained the values from a version of GPT2 fine-tuned for Dutch (‘gpt2-small-dutch’) (65) using the package *minicons* (66). Surprisal and entropy were estimated at the level of the epoch. That means that the first word of a 30-second epoch did not have any context for surprisal estimation. For surprisal, that means that the value corresponds to context-free probability. This is the same as word frequency, though estimated with a different language model. For entropy, we set the value of the first word to 0. Conversely, the last word of a segment had a context of all the preceding words that contributed to surprisal and entropy estimates.

Like the word onset and word frequency features, the surprisal and entropy features were created by inserting the values in a time-course of zeros at the samples corresponding to word onset, and the time-courses were convolved with a Gaussian window with a standard deviation of 15ms.

#### 2.5.3. Syntactic features

To create syntactic features, we required a syntactic representation of the stimulus. All sentences were manually parsed for constituency structure using a simplified minimalist framework. This entailed drawing a full X-bar structure for noun phrases (NP) and verb phrases (VP), but not for other phrases unless the intermediate projections were filled. Using the full X-bar structure for NPs and VPs ensured proper distinction between arguments and adjuncts, with arguments being attached as a sister of the head of a phrase, and adjuncts occupying the intermediate projection. See Figure 2 for the parse of an example sentence.

From the resulting structures we extracted node counts. Node counts provide an indication of the number of syntactic operations that are performed at the position of a given word. These estimates have been shown to explain variance in several types of neural recordings, such as fMRI and MEG (18,30,67,68). To count the nodes that are built for a word, we must assume a parsing algorithm. We used two algorithms: a top-down algorithm (hereafter: “top-down”) and a bottom-up algorithm (hereafter: “bottom-up”). The top-down parsing algorithm is predictive. It enumerates nodes left-to-right, building all possible nodes upward and to the right in the tree structure. This is comparable to ‘open bracket’ estimates. In other words, the top-down feature specifies the number of nodes that need to be built to incorporate future words. The bottom-up parsing algorithm, by contrast, only counts a node whenever it has seen all evidence for this node. This is comparable to ‘closing bracket’ estimates. We assumed that the parser was a ‘perfect oracle’: it only builds nodes for the correct structure when faced with ambiguity.

As can be observed in Figure 2, the syntactic analysis of a sentence may contain ‘t’. This is a *trace*, a syntactic marker that results from constituents moving to different projections in the tree structure as part of the spell-out (linearization) of the sentence. Even though they make up part of the structure, these traces have no (clear) acoustic correlate. In other words, they do not belong to any word, but they do have nodes associated with them. To solve this issue, we summed the node counts that belong to trace elements to the preceding word. As an example, consider the word ‘zeggen’ (*to say*) from the example sentence in Figure 2. The bottom-up node count for this word is 5 (upward to the left: V, VP, VP, VP VP). The bottom-up node count for the following trace is also 5 (I, IP, IP, CP, CP), leading to an assigned word count of 5+5=10 for the word ‘zeggen’.

#### 2.5.4. Model fitting and statistical analysis

Two TRF analyses were performed: (1) ‘joint encoding’, a by-electrode evaluation of the contribution of each of the experimental features (surprisal, entropy, bottom-up, and top-down); and (2) ‘interaction’, an evaluation of the influence of probabilistic properties of words on the neural encoding of syntactic structure.

##### Joint encoding

In the ‘joint encoding’ analysis, the goal was to evaluate to what extent each of the four experimental features was encoded in HGP. To this end, we estimated TRF models for all combinations of the four features of interest. As described above, each model contained the three ‘base’ features: speech envelope, word onset, and word frequency. The features in each model are summarized in Table 1 below. Estimating the models with all 16 feature combinations allowed the estimation of the contribution (or encoding) of each feature irrespective of the presence or absence of the other experimental features.

**Table 1.**
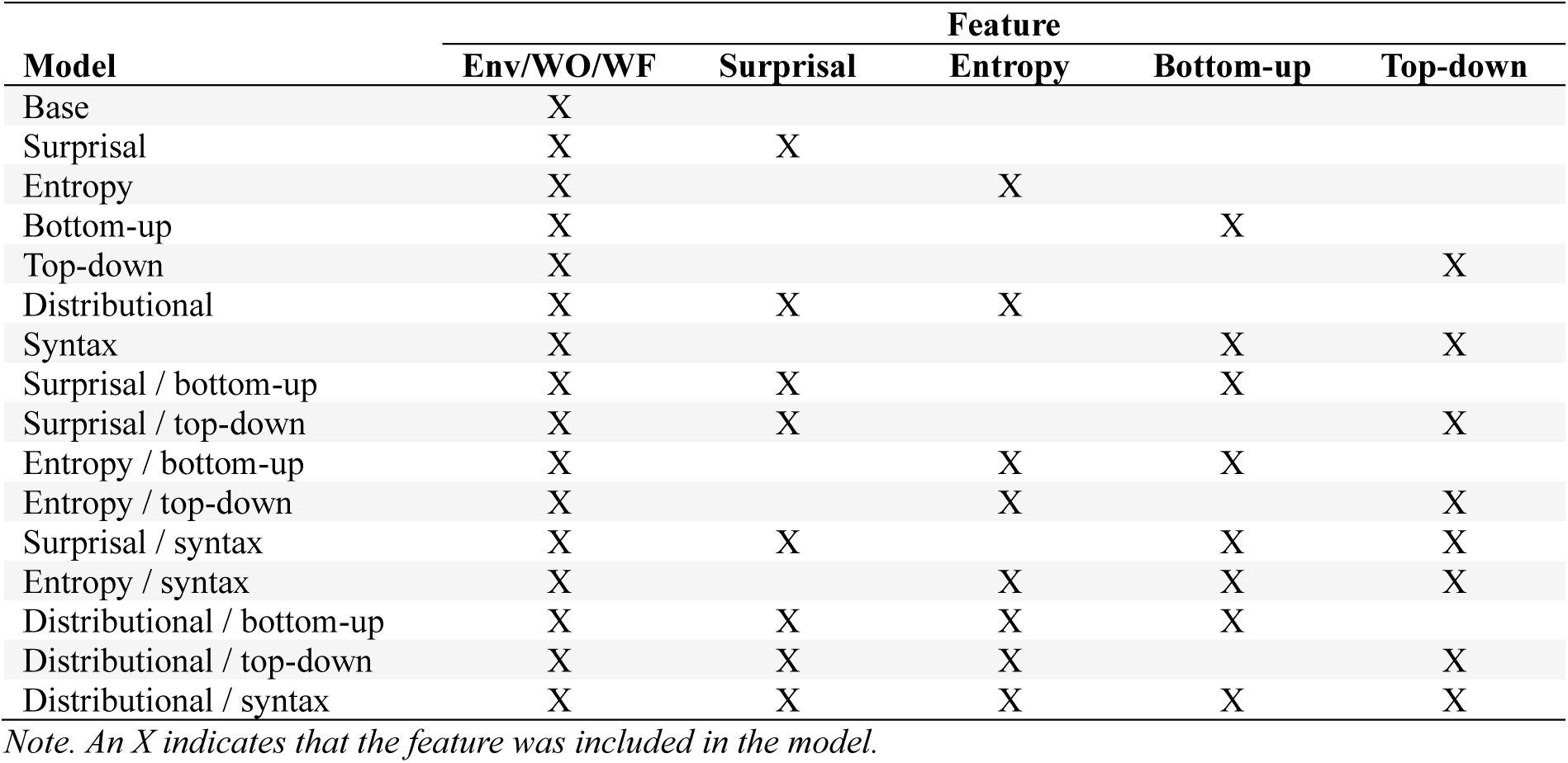
Features in each of the 16 fitted TRF models in the ‘joint encoding analysis. Abbreviations: Env. = speech envelope; WO = word onset; WF = word frequency.

We estimated the encoding of each feature in HGP by estimating the slope related to the presence or absence of the feature with a linear model applied to each electrode individually. This means that for each electrode, we created a linear model using ordinary least squares (OLS) with statsmodels (69, version 0.14.4). The predictor matrix consisted of dummy predictors that indicated the presence (1) or absence (0) of a feature in the model, like how the presence of a feature is indicated in Table 1. The dependent variable was the collection of R^2^ values for the 16 experimental models. The formula used was *R^2^ ∼ surprisal + entropy + bottomup + topdown*. Since we performed this analysis at the single-electrode level, we evaluated the significance of the coefficients by fitting 1000 OLS models with random permutations of the predictor matrix. Two-sided p-values were calculated using the observed and permuted t-values. This statistical procedure was only applied to electrodes that had a positive ‘best’ R^2^ (across models); electrodes for which the best R^2^-value was negative were skipped (N_analyzed_=554, N_skipped_ = 823; see Figure S2).

##### Interaction

The interaction analysis was conducted to study the influence of the probability of a word on the HGP correlates of syntactic structure building. Since interaction terms in TRFs do not yield interpretable waveforms and can obscure latency effects (30,70), we used a split-feature approach. This meant that we divided the syntactic node counts over two features based on the median of word probability values, namely: (Feature 1) node counts for high surprisal (or entropy), and (Feature 2) node counts for low surprisal (or entropy). This split was done for all four combinations of probability and syntactic features: bottom-up split by surprisal, bottom-up split by entropy, top-down split by surprisal, and top-down split by entropy. The evaluation of the effect of this division on reconstruction accuracy was done by fitting 1000 models with random divisions of the syntactic feature. The effect was considered significant if the ‘systematic’ split (i.e., using either surprisal or entropy) resulted in a R^2^-value that was higher than 95% (or more) of the ‘random’ split.

Like in the joint encoding analysis and to account for variation between electrodes, we fit each division with various combinations of additional features. All feature combinations are summarized in Table 2 below. We used the results of the ‘main effects’ analysis to determine which model to evaluate and interpret for each electrode. Specifically, we checked for each electrode which features had positive coefficients for their contribution to the signal (irrespective of their significance). We then selected the model that contained all these features and one of the splits. For example, if an electrode had non-negative effects for surprisal, entropy, and bottom-up, we would evaluate the models that split bottom-up by either surprisal or entropy, and also contain surprisal and entropy as main effects (i.e., rows 11 and 12 from Table 2 below) against their random counterparts.

**Table 2.** Features in each of the 16 fitted TRF models in the ‘split feature’ analysis. Abbreviations: Env. = speech envelope; WO = word onset; WF = word frequency; surp. = surprisal; entr. = entropy; BU = bottom-up; TD = top-down.

| Model | Feature |  |  |  |  |  |  |  |  |
| --- | --- | --- | --- | --- | --- | --- | --- | --- | --- |
|  | Env/<br>WO/<br>WF | Surp. | Entr. | BU | TD | BU<br>split<br>by<br>surp. | BU<br>split<br>by<br>entr. | TD<br>split<br>by<br>surp. | TD<br>split<br>by<br>entr. |
| Surprisal/top-down | X | X |  |  |  |  |  | X |  |
| Surprisal/bottom-up | X | X |  |  |  | X |  |  |  |
| Surprisal/bottom-up + top-down | X | X |  |  | X | X |  |  |  |
| Surprisal/top-down + bottom-up | X | X |  | X |  |  |  | X |  |
| Entropy/top-down | X |  | X |  |  |  |  |  | X |
| Entropy/bottom-up | X |  | X |  |  |  | X |  |  |
| Entropy/bottom-up + top-down | X |  | X |  | X |  | X |  |  |
| Entropy/top-down + bottom-up | X |  | X | X |  |  |  |  | X |
| Surprisal/top-down + distributional | X | X | X |  |  |  |  | X |  |
| Entropy/top-down + distributional | X | X | X |  |  |  |  |  | X |
| Surprisal/bottom-up + distributional | X | X | X |  |  | X |  |  |  |
| Entropy/bottom-up + distributional | X | X | X |  |  |  | X |  |  |
| Surprisal/top-down + distributional + bottom-up | X | X | X | X |  |  |  | X |  |
| Entropy/top-down + distributional + bottom-up | X | X | X | X |  |  |  |  | X |
| Surprisal/bottom-up + distributional + top-down | X | X | X |  | X | X |  |  |  |
| Entropy/bottom-up + distributional + top-down | X | X | X |  | X |  | X |  |  |

Note. An X indicates that the feature was included in the model.

If the interaction resulted significant from the permutation test on the R^2^-values, we evaluated the difference between the TRF waveforms. For this, we computed the difference between the ‘high probability’ and ‘low probability’ waveforms at each time-point. We did the same for the 1000 random divisions. We marked as significant the time-points for which the difference between the systematic split was in the upper 97.5% or lower 2.5% of the random distribution.

## 3. Results

### 3.1. Joint encoding

#### 3.1.1. Reconstruction accuracy

In the ‘joint encoding’ analysis, the goal was to evaluate to what extent each of the four experimental features was encoded in HGP, and whether (and how) multiple features are encoded in any electrodes. We fit TRF models with all possible combinations of our experimental features – surprisal, entropy, bottom-up node counts, and top-down node counts. Reconstruction accuracy values for the smallest and the largest models (‘base’ and ‘distributional/syntax’, respectively, see table 1 above) and the raw difference (‘distributional/syntax’ – ‘base’) between these are shown in Figure 3 below. As is apparent from the first two panels of this plot, reconstruction accuracy was qualitatively highest for electrodes along the sylvian fissure and some electrodes in the middle temporal gyrus. The third panel in Figure 3 indicates that adding all four features to the base model numerically improves the reconstruction accuracy in several electrodes in this same area, with smaller increases further away from the sylvian fissure.

**Figure 3.**
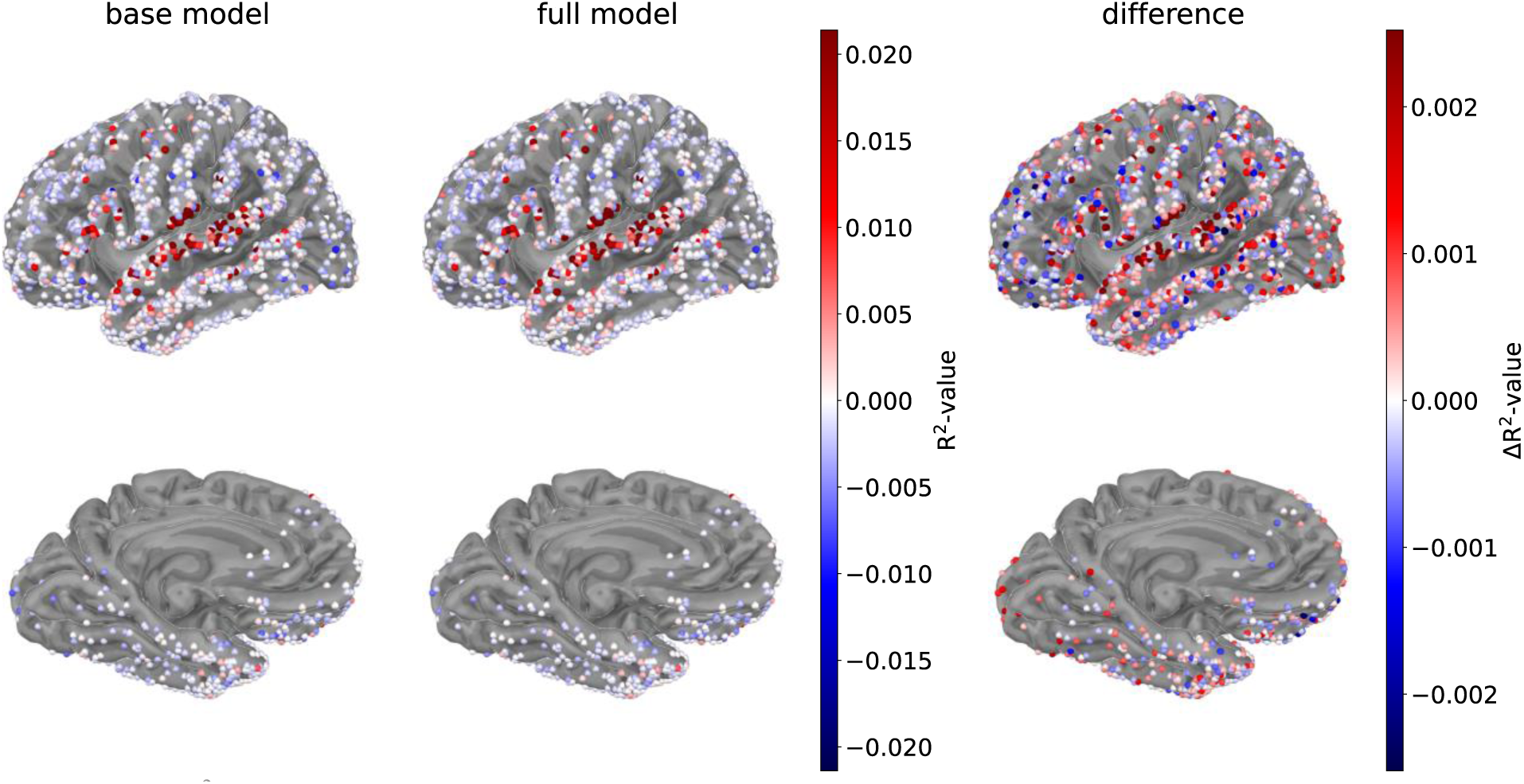
Raw R2-values per electrode for the base model (speech envelope, word onsets, and word frequency) and the largest model (speech envelope, word onsets, word frequency, surprisal, entropy, bottom-up and top-down). The rightmost panel shows their difference by subtracting the base model from the largest model. N.B. The largest model is not necessarily the best model. The figure displays the left hemispheric pial surface of the fsaverage brain. The medial view is tilted by 130° to visualize the ventral temporal electrodes.

Prior to statistical analysis, we removed the electrodes in which R^2^-value of the best-fitting model (among the models in Table 1) was negative; 554 electrodes were retained (see Figure S2). As described in 2.5.4. ‘Joint encoding’, the contribution of each experimental feature (surprisal, entropy, bottom-up, top-down) to model fit was evaluated by entering the R^2^-values of all 16 models into OLS models with predictors indicating the presence (1) or absence (0) of each feature. This analysis provides, for each electrode and for each feature, a coefficient (the estimated contribution of the feature to the R^2^-value; Figure 4) and a t-value (capturing the statistical reliability of the coefficient; corresponds to the coefficient divided by its standard error; Figure 5). Significance of the coefficients was evaluated using permutation testing (Figure 4, Figure 5).

**Figure 4.**
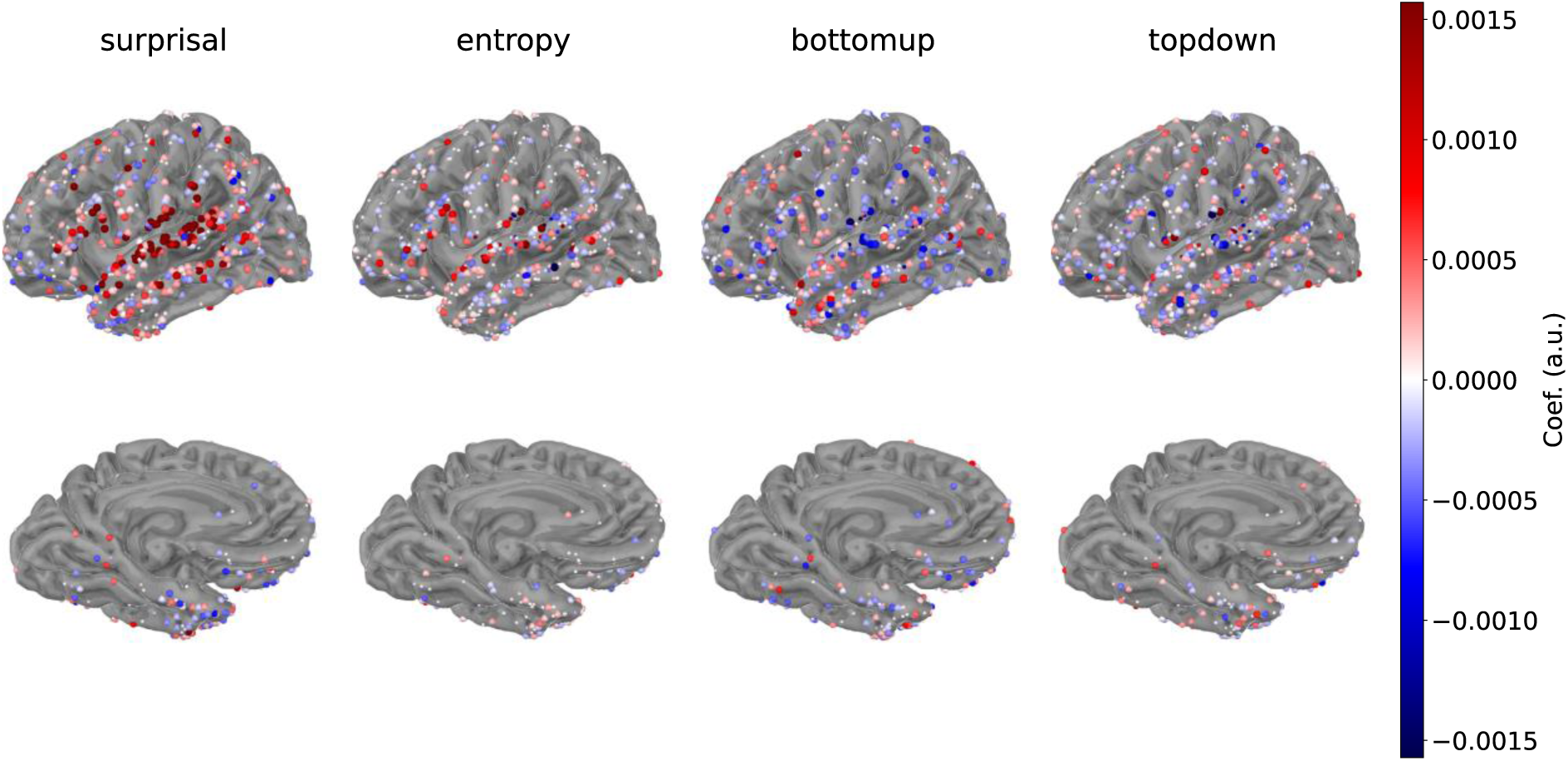
The coefficients that describe the contribution of each feature to the model of the signal. Blue indicates a negative contribution to a model of the signal; red indicates a positive contribution to a model of the signal. Smaller (mostly white) electrodes were insignificant. The figure displays the left hemispheric pial surface of the fsaverage brain. The medial view is tilted by 130° to visualize the ventral temporal electrodes.

**Figure 5.**
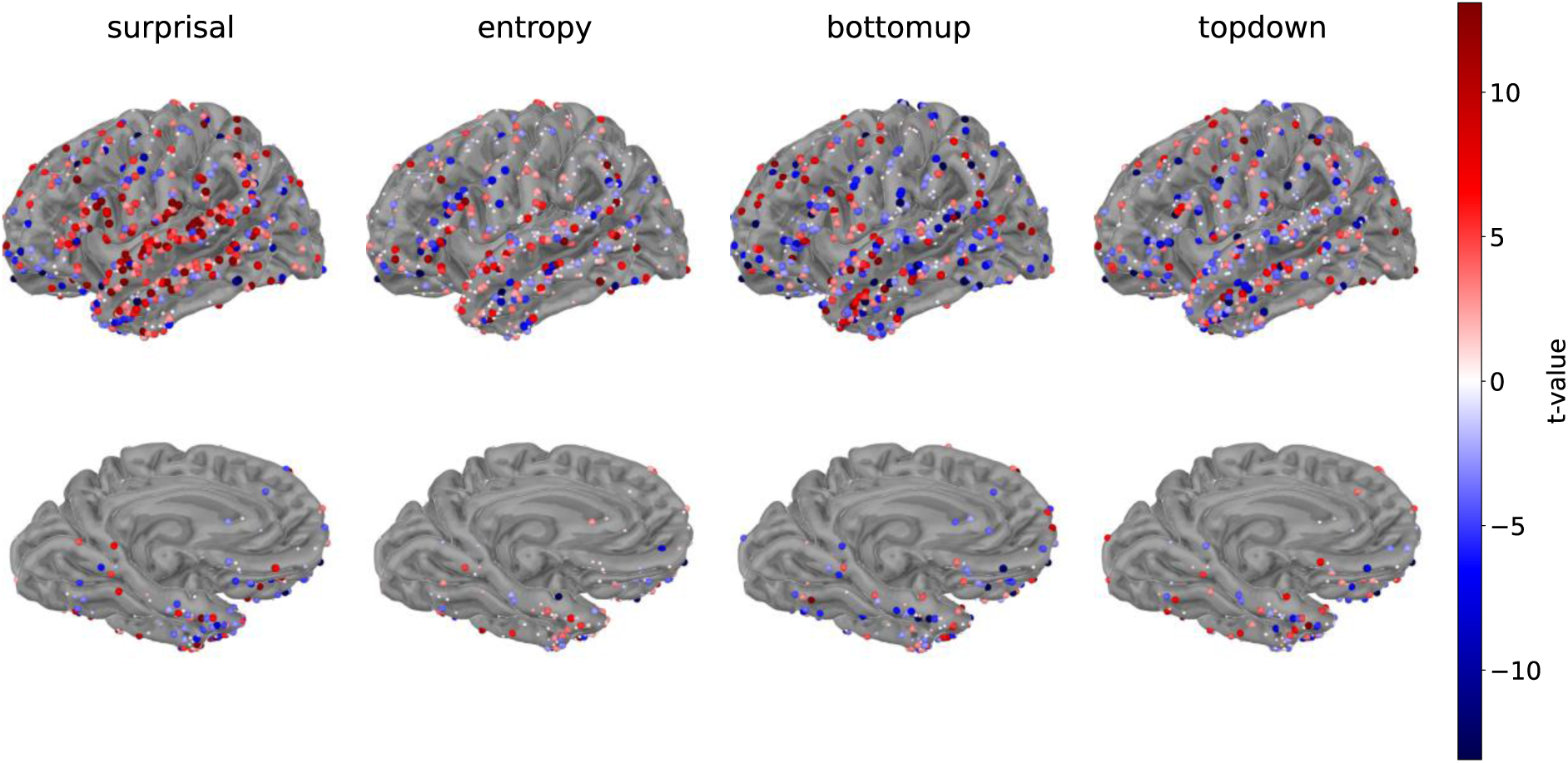
T-values for the coefficients that describe the contribution of each feature to the model of the signal. Blue indicates a negative contribution to a model of the signal; red indicates a positive contribution to a model of the signal. Smaller (mostly white) electrodes were insignificant. The figure displays the left hemispheric pial surface of the fsaverage brain. The medial view is tilted by 130° to visualize the ventral temporal electrodes.

The analyses revealed that all four features had a significant contribution to models of the signal, though their contributions differed in magnitude and in their spatial distribution. As shown in Figure 4, surprisal made relatively the largest contribution to the R^2^-values, both in terms of the number of electrodes and in terms of coefficient values. The largest coefficients were observed in the STG and the pre-and postcentral operculum, as well as a few electrodes in pars triangularis and pars opercularis of the IFG. Figure 5 illustrates the higher statistical reliability of these coefficients. Smaller effects are found in the MTG and supramarginal gyrus (SMG). Coefficients for entropy were smaller than for surprisal, which contributed positively in STG, with its distribution being more anterior than that of surprisal, covering the temporal pole and the anterior ventral portion of the temporal lobe. Entropy also contributed to models of electrodes in the entire IFG; though for this feature, electrodes with a positive coefficient were located directly alongside electrodes with negative coefficients. This suggests that there is more variability in the spatial distribution of the encoding of entropy than for surprisal.

The significantly positive coefficients for structure building (bottom-up and top-down node counts) were smaller and less widespread (Figure 4), but their effects were reliable (Figure 5). The strongest contributions of bottom-up node counts appeared in anterior areas, located in pars triangularis, the anterior portion of the STG and MTG, and a group of electrodes in the dorsolateral prefrontal cortex. In stark contrast to surprisal, adding bottom-up node counts *decreased* the reconstruction accuracy of models in the posterior part of the superior- and middle temporal gyrus, suggesting that the feature is not encoded in these areas. The contributions of top-down node counts appear distributed, with a row of positive contributions in the SMG and inferior parietal cortex, a few electrodes in pars orbitalis, and some clusters in the MTG. For comparison, we provide the contrasts of raw R^2^- values between the base model and base plus each individual feature, and between the full model and full minus each individual feature in supplementary figures S3 and S4.

Having seen that all features contribute to HGP, we evaluated their joint encoding: to what extent do features co-occur in the same electrodes? We evaluated this in two ways. Firstly, we visualized the combinations of significant features per electrode; and, secondly, we estimated the correlations between feature coefficients.

Figure 6 provides an overview of all (significant) combinations of features. The encoding of a single feature indicates the existence of neural populations that selectively encode this feature. Joint encoding, on the other hand, could arise either if neural populations encoding distinct features are so close together that one ECoG electrode records activity from both (i.e., within a radius of 5 mm), or if there exist neural populations that encode both types of information. The data at hand cannot adjudicate between these two possibilities.

**Figure 6.**
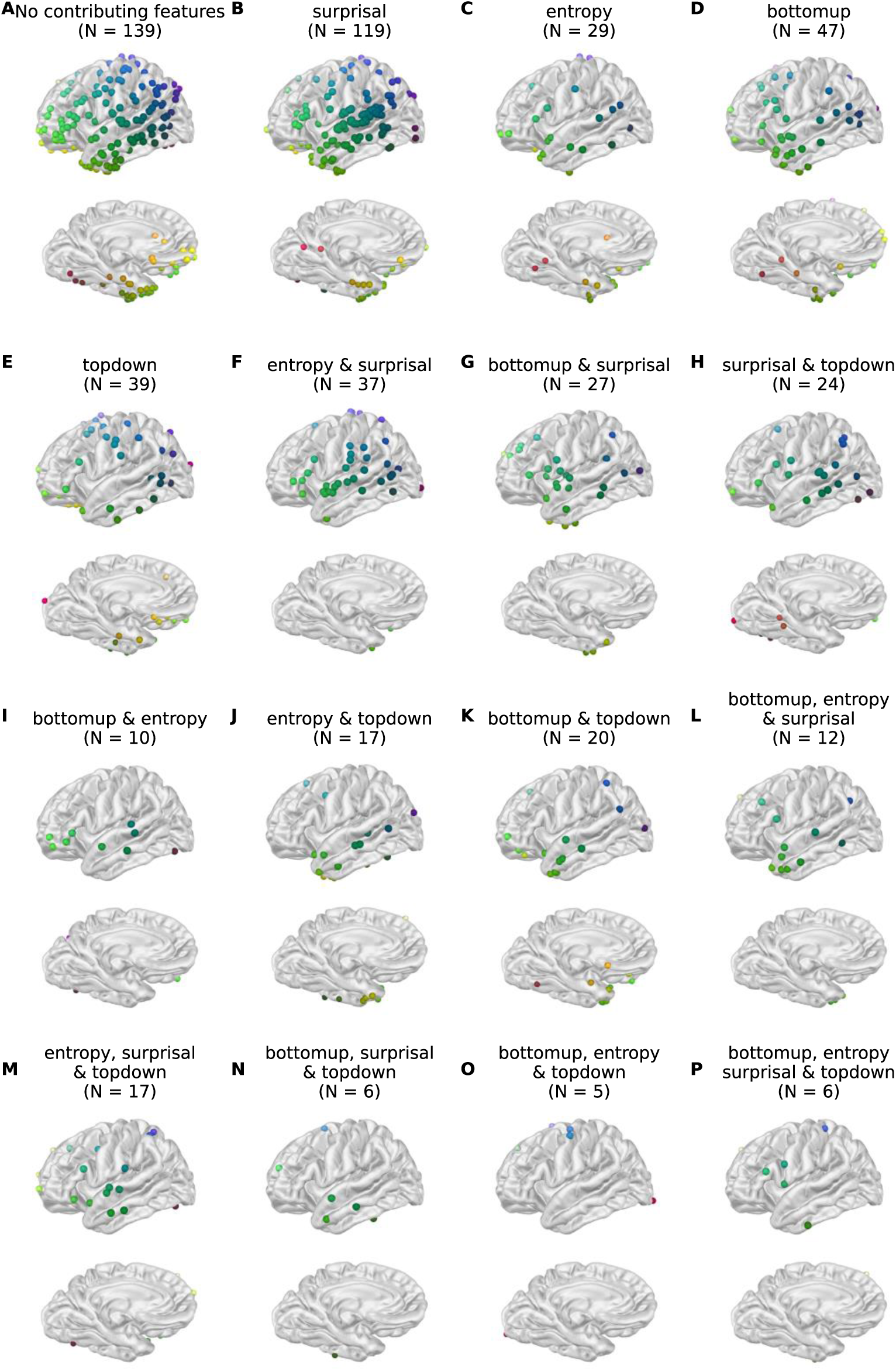
Sensitive electrodes to each feature and feature combinations. A electrode is displayed if the feature (or features) in question (all) made a significantly positive contribution to the model of HGP, while no other features did. In other words, the electrodes displayed at the plot ‘surprisal’ had significant positive coefficients for surprisal and only surprisal; the electrodes displayed at ‘entropy & surprisal’ had significant positive effects of both entropy and surprisal, and no other features. The figure displays the left hemispheric pial surface of the fsaverage brain.The medial view is tilted by 130° to visualize the ventral temporal electrodes. (A) No contributing features. (B). Surprisal. (C) Entropy. (D) Bottom-up. (E) Top-down. (F) Entropy and surprisal. (G) Bottom-up and surprisal. (H) Surprisal and top-down. (I) Bottom-up and entropy. (J) Entropy and top-down. (K) Bottom-up and top-down. (L) Bottom-up, entropy, and surprisal. (M) Entropy, surprisal, and top-down. (N) Bottom-up, surprisal, and top-down. (O) Bottom-up, entropy, and top-down. (P) Bottom-up, entropy, surprisal, and top-down (all features).

We observed that all possible combinations existed: the dataset included electrodes that encode all features (N=6), each sub-combination of features, and all features individually. In 139 of the 554 analyzed electrodes, none of the features explained additional variance as compared to the base model. 234 electrodes were sensitive to only one feature, and the remaining 181 encoded various combinations of features. Surprisal was encoded in the highest number of electrodes: in 119 electrodes it was the only encoded feature, and in 129 other electrodes it occurred jointly with other features. The electrodes that encode only surprisal form a cluster in the posterior STG, and another one in pars triangularis. In 29 electrodes, entropy was the only encoded feature. Entropy co-occurred with other features in 104 electrodes. Bottom-up was the only encoded feature in 47 electrodes. These electrodes are divided over the anterior and posterior portion of the temporal lobe and the prefrontal cortex, with a small cluster in pars orbitalis and the dorsolateral prefrontal cortex. Bottom-up co-occurred with other features in 86 electrodes. Top-down was the only encoded feature in 39 electrodes. These electrodes appeared dispersed and more distant from the perisylvian fissure than those of the other features: in the posterior MTG and inferior temporal gyrus (ITG), pre-and postcentral gyrus, dorsolateral prefrontal gyrus. Top-down co-occurred with other features in 95 electrodes. In total, 106 electrodes were selectively sensitive to syntax; 185 were selectively sensitive to probability; and 124 encoded both types of features. As shown in Figure 7, the spatial distribution of the feature types is similar; both effects are widespread, though very few electrodes in the middle MTG and STG and posterior STG appear to encode a structural feature alone.

**Figure 7.**
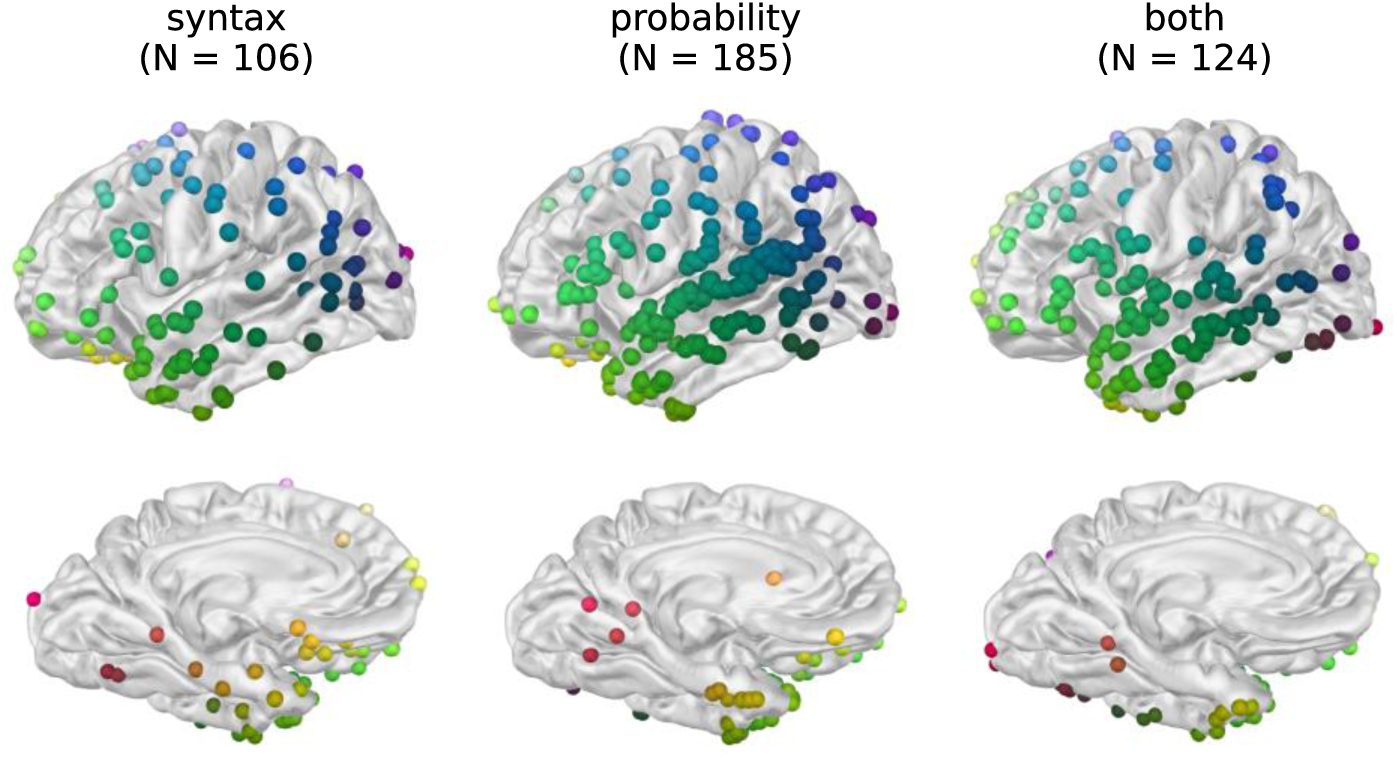
Sensitive electrodes grouped by feature type. ‘Syntax’ electrodes had significant, positive effects for bottom-up node counts, top-down node counts, or both. ‘Probability’ electrodes had significant, positive effects for entropy and surprisal. ‘Both’ electrodes had positive effects for one or more features of each feature type. RGB colors indicate position. The figure displays the left hemispheric pial surface of the fsaverage brain. The medial view is tilted by 130° to visualize the ventral temporal electrodes.

Subsequently, we computed the Pearson’s R correlation matrix between all feature coefficients across the (N=554) electrodes. A positive correlation between the coefficients of two features indicates that the contributions of these features similarly vary across electrodes, suggesting that they are encoded by shared neural substrate; while a negative correlation indicates that features are encoded by separable neural populations: neural populations that are sensitive to one feature, are less likely to be sensitive to the other. This approach quantifies the degree of joint encoding in a continuous fashion. Given non-normality of the distributions – most notably surprisal; see Figure S5 –, the coefficients were Yeo-Johnson transformed prior to the correlation analysis.

As displayed in Figure 8, this analysis revealed that the coefficients for bottom-up node counts moderately negatively correlate with both surprisal (r = -.15, p < .01) and entropy (r = -.13, p < .01), suggesting that coefficients for bottom-up node counts decrease as the coefficients for lexical probability increase. In other words, if an electrode strongly encodes entropy or surprisal, the contribution of bottom-up node counts to model fit is more likely to be small or even negative. This is visible particularly in the posterior STG as shown in Figures 5 and 6, where the coefficients for surprisal and entropy are largely positive, while those for bottom-up are mostly negative. This suggests that the neural substrates for bottom-up node counts and lexical probabilistic information are (partially) separable.

**Figure 8.**
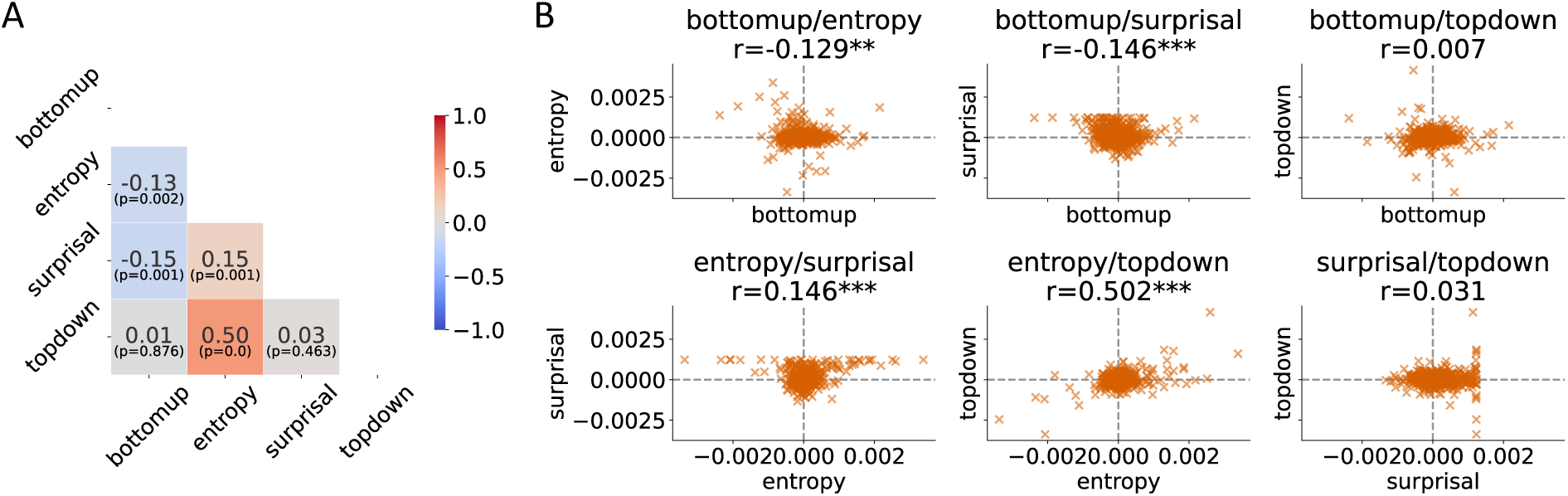
Pearson’s R correlation between the coefficients for each feature. (A) The correlation matrix including Pearson’s R correlation coefficient and its significance. (B) Scatterplots for all feature combinations. Each ‘x’ is one electrode. All features were individually Yeo-Johnson transformed for the analysis and display in this figure. P-value indicators: <= .05 *; < .01 **; < .001 ***

For top-down node counts, on the other hand, there was a strong positive correlation with entropy (r = .50, p < .01), which indicates that electrodes that have larger (or smaller) coefficients for entropy also tend to have larger (or smaller) coefficients for top-down. This shared variability suggests that entropy and top-down are (partially) encoded by shared neural populations. There was also a positive correlation between entropy and surprisal (r = .15, p < .01), suggesting that surprisal and entropy are (partially) encoded by shared neural populations.

#### 3.1.2. Time courses

We then examined the TRFs – i.e., the time courses of beta weights – for each feature, in the electrodes that encoded this feature significantly as identified based on the reconstruction accuracy values – i.e., the electrodes depicted large and in red in Figures 5 and 6. This tells us not only when effects are happening, but also their direction: does HGP increase or decrease as a function of a feature? To identify global patterns in the time-courses, we smoothed the TRF time courses by convolving them with a gaussian window with a standard deviation of 50ms, and subsequentially identified peaks for each electrode using SciPy’s peak detection algorithm. Only peaks with the 30% highest prominence were retained. Using an adjacency matrix across all electrodes projected onto the fsaverage template (electrodes at the 10^th^ percentile of the distance distribution or less were considered adjacent), we identified spatiotemporal clusters by finding groups of minimally 5 adjacent electrodes of which the signal peaked within 100ms surrounding the original peak. These clusters are plotted as bold lines in the plots in Figure 9 below. The unfiltered HGP TRF time courses are provided in Figure S6; videos of the time-courses can be accessed here: https://osf.io/vf2dg/.

**Figure 9.**
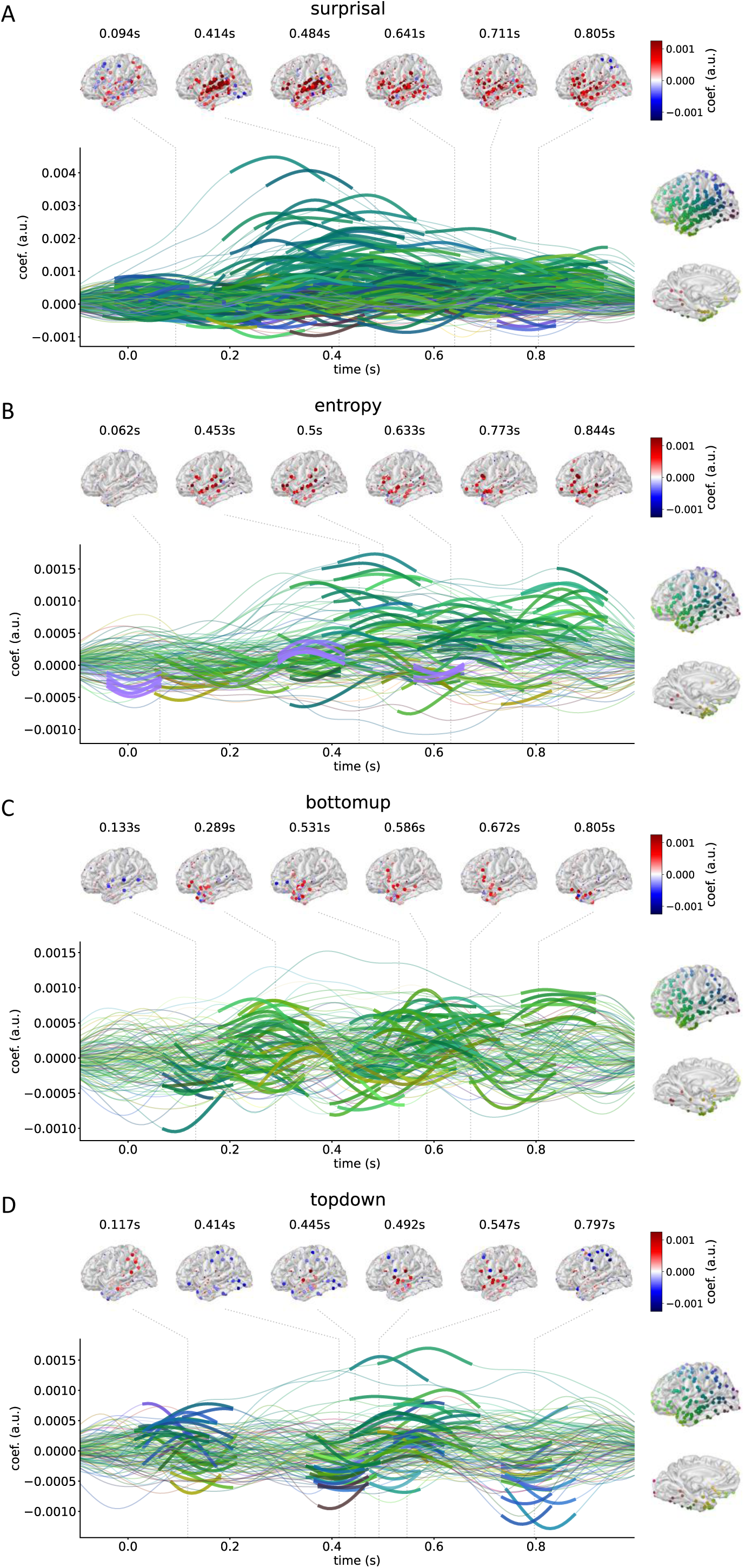
Temporal response functions for the four features. (A) Surprisal; (B) entropy; (C) bottom-up; (D) top-down. The displayed electrodes are those that had significant, positive coefficients for the feature in question. Lines printed in bold were identified using a clustering method that identifies peaks in 5 adjacent electrodes within a timespan of 100ms. RGB colors indicate electrode position. The row of brain maps shows the spatial distribution of the effects. Large electrodes are part of a cluster. The timepoints of the brain maps were identified using the maximum absolute amplitude of all the electrodes and time-points that were part of a cluster. To display the variation in all features, the temporal response function for surprisal was plotted with a wider scale on the y-axis.

As can be observed in Figure 9A, the surprisal TRF revealed a single large, positive response indicating that HGP increases as a function of increasing surprisal. The pattern is as follows. The TRF amplitude peaks between 300 and 500 milliseconds depending on the electrodes, with the largest coefficients in the posterior STG. Across the whole time-window, the shift towards strongly positive coefficients spreads spatially. The response appears to start with moderately positive coefficients around 100 milliseconds in the anterior temporal lobe, and to increase across time. By 300 milliseconds, HGP peaks in the STG, and increases in SMG and the operculum. Even though the total amplitude decreases after approximately 400 milliseconds, the spatial spread continues, and by 800 milliseconds the whole IFG, temporal pole, and some electrodes in the dorsolateral prefrontal cortex see an increase of HGP as a function of surprisal. This suggests that HGP increases as words become less probable given the context, and that this effect spreads from temporal areas to frontal areas over 900 milliseconds. There are a few electrodes that suggest negative coefficients, though these are of smaller magnitude than the positive coefficients. The entropy TRF similarly revealed one large response that starts around 200 milliseconds and peaks around 500 milliseconds in the anterior STG, MTG, and IFG. Besides this pattern, a group of electrodes coming from an HD grid placed on the lateral pre- and postcentral gyrus (one participant; the purple lines in Figure 9B) indicates an early response, decreasing as a function of increasing entropy around word onset and increasing between 350 and 400 milliseconds. Similarly to surprisal, some electrodes are associated with negative coefficients. These results indicate that HGP increases as a function of both increasing uncertainty (entropy) and surprise (surprisal) in most electrodes. For both features, some electrodes revealed negative coefficients. The effects for surprisal and entropy are both largest around 400-500 milliseconds and in the STG/IFG, though generally the effect of entropy appears slightly later and slightly more anterior than the effect of surprisal.

While the lexical probabilistic features suggest single-stage responses, the picture is different for the syntactic features. The response to bottom-up node counts (Figure 9C) is most apparent for electrodes in pars triangularis, pars opercularis, and the anterior portion of the temporal lobe. The effect of bottom-up node count on cortical activity in these areas suggests a multi-phasic response – an ‘early’ one, around 300 milliseconds, and a late one, around 600 milliseconds. To be precise, coefficients are close to zero or slightly negative in the first 150 milliseconds. HGP shifts from decreasing to increasing as a function of increased bottom-up node count in

IFG and the temporal pole around 290 milliseconds, decreases around 400 milliseconds, and increases again between 500 and 700 milliseconds in the same area. The electrodes in the temporal pole then suggest another simultaneous in- and decrease of HGP around 800 milliseconds, while electrodes in IFG and the anterior STG remain positive. For top-down node counts (Figure 9D), we also observed a multi-stage pattern that involved electrodes in the STG, the parietal cortex and electrodes along the central sulcus. This started with a brief increase of HGP in the parietal cortex and SMG around 100 milliseconds, followed by a decrease in HGP in areas further away from the sylvian fissure and an increase in middle-to-anterior STG and MTG. Then, approximately 800 milliseconds post word onset, HGP decreases as a function of top-down node count in the electrodes in parietal cortex and around the central sulcus. These results suggest that structure building is not an instantaneous process; rather, it suggests that structure building is a process that requires intermittent communication between two or more (distant) areas.

#### 3.1.3. Joint encoding - summary

In summary, the results from the joint encoding analysis suggest that surprisal, entropy, bottom-up node counts and top-down node counts are all encoded by HGP, both in isolation and jointly. Surprisal was encoded in most of the recorded signals and had the largest contribution in terms of coefficients. The effects are largest in the posterior STG and pars triangularis and opercularis of the IFG. Entropy was encoded in the STG and IFG as well, though its coefficients were smaller than for surprisal and the positive coefficients had a more anterior distribution. Bottom-up node counts had positive effects in pars triangularis, the anterior portion of the STG and MTG, and electrodes in the dorsolateral prefrontal cortex. The positive contributions of top-down node counts were distributed, among others in the SMG and inferior parietal cortex, a few electrodes in pars orbitalis and the MTG. While all possible combinations of features exist in the data, correlations between feature coefficients further indicated that the contribution from bottom-up node counts is smaller when entropy and surprisal explain more variance, but that top-down is more likely to have a greater impact when entropy does, too. This suggests that top-down syntactic representations might share neural resources with entropy, while bottom-up syntactic representations are more likely to be encoded by different neural populations than lexical probability.

A clustering approach of the TRFs showed that overall, HGP globally increases as a function of both increasing uncertainty (entropy) and surprise (surprisal). The peaks are around 400-500 milliseconds in the STG/IFG, though generally the effect of entropy appears slightly later and slightly more anterior than the effect of surprisal. The TRFs for structure building suggest multi-stage approaches in which HGP increases and decreases alternatingly, which recruit IFG and the anterior temporal lobe (bottom-up) or parietal areas (top-down). These results suggest that structure building is not an instantaneous process; rather, it suggests that structure building is a process that requires intermittent communication between two or more (distant) areas.

### 3.2. Interaction analysis

#### 3.2.1. Reconstruction accuracy

Having shown that lexical and syntactic features may be encoded within the same, or neighbouring populations, as reflected in the patterns of results described above, we sought evidence for interaction between these. In line with suggestions from the statistical learning literature (27,28), we examined whether lexical probability affected cortical activity related to structure building. To this end, we tested four models in which node counts were divided according to the value of the lexical probabilistic features: top-down node counts divided by the median of surprisal or entropy; and bottom-up node counts divided by surprisal or entropy. In other words, there were two syntactic features in each model: one with the node counts of words for which the surprisal or entropy value was above the median, and with the node counts of the words for which the surprisal or entropy value was below the median. The reconstruction accuracy of these models was compared against a distribution of 1000 models with node counts randomly divided over two features. If splitting according to the probability values results in a better fit than splitting randomly, we can conclude that there is some effect of the probability feature on the encoding of the syntactic feature: an interaction.

Multiple electrodes showed an interaction between bottom-up node counts or top-down node counts by surprisal (Figure 10). The interaction between surprisal and bottom-up node count was found in 14 electrodes, of which six were located along the STG, one in pars opercularis, two in pars triangularis, two in the SMG, one in the posterior operculum, and two in the dorsolateral prefrontal cortex. The interaction between top-down node count and surprisal was found in 14 electrodes, with six electrodes on the STG, two in the MTG, one in the SMG, one in the posterior operculum, one in the precentral gyrus, two in the dorsolateral prefrontal cortex, and one in pars triangularis. Interactions between syntactic features and entropy were less apparent. An effect of splitting bottom-up node count by entropy was found in six electrodes. Two of these were in the IFG – pars opercularis/triangularis –, one in the anterior MTG, one in the anterior ITG, and one on the precentral gyrus. Splitting by entropy had a significant effect on responses to top-down node counts in 11 neuroanatomically dispersed electrodes: two were in pars opercularis, two on the ventral side of the temporal pole, one in the posterior operculum, one in the parahippocampal gyrus, and two in the dorsolateral prefrontal cortex. Despite the small number of electrodes, these results suggest that the process of structure building is mediated by lexical probability in multiple ways.

**Figure 10.**
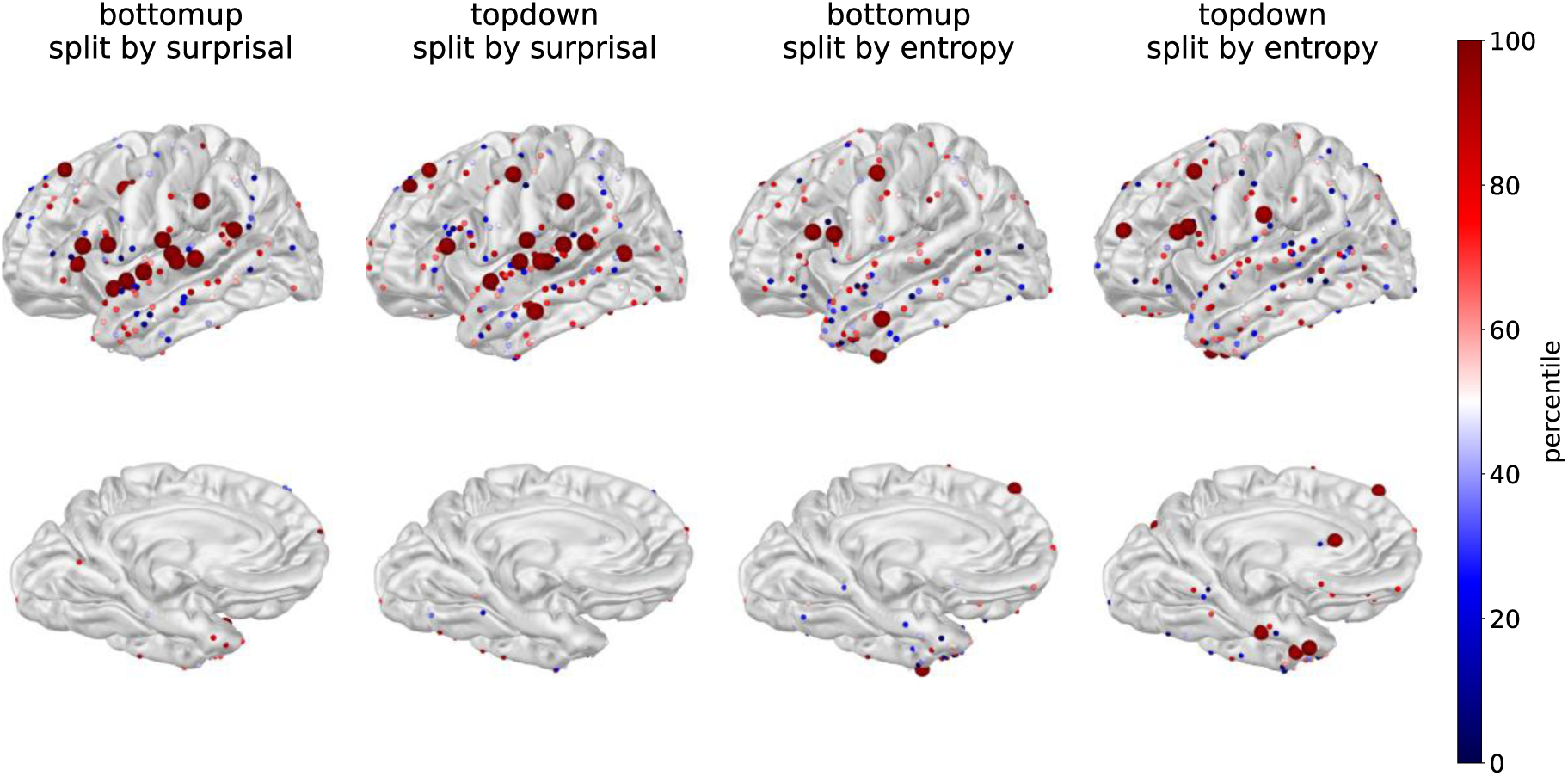
Spatial distribution of the results from the split features-analysis. The coloring indicates the percentile of the R^2^-value of the model with a feature that was systematically split (by entropy or surprisal) on the distribution of R^2^-values of1000 models in which the feature was split randomly. The larger electrodes were significant – i.e., had R^2^-values that sat at 95% of the random distribution.

#### 3.2.2. Time courses

The (temporal) differences between the TRFs for the ‘high’ and ‘low’ subsets of the data were determined by comparing the difference between the split-feature TRFs with the difference between the TRFs from the random split models. This revealed a pattern for the effect of surprisal on the syntactic features: high surprisal words are associated with more positive TRF-coefficients than low surprisal words, suggesting that HGP increases more with increasing node counts if a word has low predictability. This holds for both top-down and bottom-up node counts.

For bottom-up node counts, the effect of surprisal (Figure 11A) appears driven by coefficients being more positive when surprisal is high, and for some electrodes even negative when surprisal is low. The effect appears reversed – i.e., more negative for high surprisal words – for one electrode in the posterior STG. Because the ‘high surprisal’ variant of the bottom-up TRF visually most resembled the TRF estimated on the whole dataset of the same electrodes (provided in the leftmost column of Figure 11 below), we correlated the ‘original’ TRF with both the low-surprisal TRF and the high-surprisal TRF (before smoothing). While the correlations were positive and significant in both conditions (mean r_high_ = .74 (sd .09), mean r_low_ = .60 (sd .18); all p <.01), the correlations between the original bottom-up TRF and the high-surprisal variants were higher than those with the low-surprisal variants (t(13)=2.70, p < .05)). This suggests that the effects of bottom-up node counts are driven mainly by high surprisal words.

**Figure 11.**
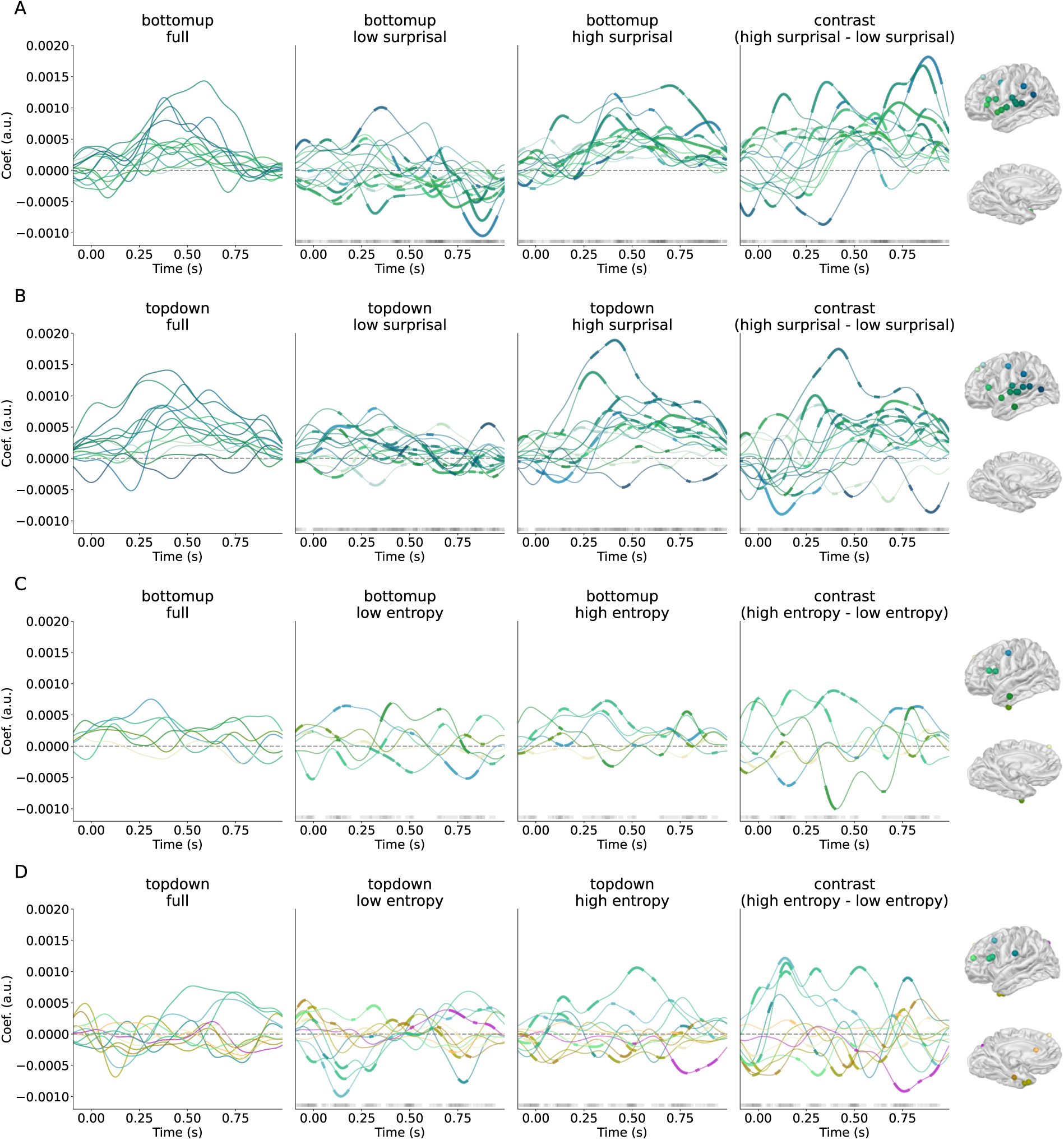
Temporal Response Functions for the split features smoothed with a Gaussian window with a standard deviation of 50 ms. (A) Bottom-up split by surprisal; (B) Top-down split by surprisal; (C) bottom-up split by entropy; (D) top-down split by entropy. Only time courses for significant electrodes are displayed. The first column shows the TRF for the feature trained on all the data (no split) for the same electrodes. The middle two columns show the halves of the split feature. The last column displays the contrast. Electrodes and time-points in bold were those that were in the upper- or lower 2.5% of the distribution of the difference between 1000 random splits. The grey bar at the bottom shows the time-points; darker grey means more electrodes were significant at that time-point. RGB colors of lines in plots reflect electrode location as shown projected on canonical brain surface at right.

For top-down, the positive difference (the rightmost column in Figure 11B) has the largest amplitude between 200 and 500 milliseconds in two electrodes in the posterior STG. Two electrodes show a negative effect, with smaller coefficients for high surprisal words than for low surprisal words. This effect is early for the electrode in the precentral gyrus (between word-onset and 250 milliseconds) and late for the electrode in the posterior MTG (250+ milliseconds). While the high- and low surprisal TRFs all significantly correlate with the original (mean r_high_ = .62 (sd .19), mean r_low_ = .60 (sd .12); all p < .01), there is no difference between the correlations of the high- and low surprisal TRFs for top-down with the ‘original’ TRF for top-down (t(13)=0.31, p = .76)), which indicates that the difference is one of amplitude alone.

The effects of entropy on syntactic structure building are more widely spatially distributed, and the effects do not appear to cluster around a specific time-point. For the effect of entropy on bottom-up, both the high- and low entropy TRFs correlate with the original bottom-up TRF (mean r_high_ = .59 (sd .09), mean r_low_ = .62 (sd .10), all p < .01), and there is no difference between the correlations of the high- and low entropy TRFs and the original bottom-up TRF (t(5)=0.60, p = .58)). For the effect of entropy on top-down, all correlations between the original top-down TRF and the high/low entropy split TRFs were significant (mean r_high_ = .55 (sd .11), mean r_low_ = .69 (sd .10), all p < .01). There was a significant difference between the correlations (t(10)=-2.39, p < .05), indicating that the low entropy TRF for top-down resembles the original top-down TRF more than the high-entropy TRF. This suggests that, in these electrodes, the time-course of top-down structure building is shaped by low-entropy words more than by high-entropy words.

#### 3.2.3. Split features - summary

Evidence was found for all four tested interactions between lexical and syntactic features: bottom-up interacted with surprisal in 14 electrodes; top-down interacted with surprisal in 14 electrodes; bottom-up interacted with entropy in 5 electrodes; and 11 electrodes revealed an interaction between top-down and entropy. Post hoc pairwise contrasts of the TRF waveforms suggested the effects of bottom-up node counts are driven mainly by high surprisal words. By contrast, the TRF of top-down node counts appeared to be shaped by low-entropy words more than by high entropy words. Despite the low number of electrodes identified here, these results suggest that lexical probability and syntactic structure actively interact in HGP. More specifically, the results indicate that lexical probability can shape the process of structure building.

## 4. Discussion

In this study, we investigated whether high gamma power (HGP) jointly encodes lexical probability and syntactic structure; and whether lexical probability affects HGP signatures of structure building. We analyzed ECoG recordings of naturalistic speech processing using temporal response functions (TRFs) in two separate analyses, testing the independent and dependent encoding of lexical and syntactic features respectively. We demonstrate that lexical probability and syntactic structure are encoded by HGP, in both spatially separate (one feature per electrode) and spatially intersecting (multiple features per electrode) fashion. In addition, a correlation analysis suggested that features may (or may not) co-occur systematically. This means that HGP indeed jointly represents probability and syntactic structure. We also find evidence for interactions between the encoding of probabilistic and syntactic information, indicating an impact of lexical probability on processes associated with structure building. With this, the current study contributes to a growing literature that suggests that lexical probabilistic and syntactic cues are exploited flexibly during speech comprehension (1,2,71–73).

### 4.1. High gamma power encodes constituency structure

Both bottom-up and top-down node counts contributed positively to the reconstruction accuracy (R^2^) of TRF models of HGP, over and above any effects of speech envelope, word onset, word frequency, surprisal, and entropy. This suggests that HGP reflects the processing of constituency structure. Previous studies have reported mixed results. While encoding of multi-unit syntactic node opening and closing measures in HGP has been found (43), other authors have suggested that the effect can be accounted for by including word frequency as a predictor (44). Consequently, Murphy (9) proposed that HGP captures basic semantic composition (3), but is not sensitive to multi-unit syntactic node opening and closure. Because the present study shows effects of top-down and bottom-up node counts derived from constituency structure over and above word frequency, the results reported here are not consistent with this interpretation. This may be due to differences in parsing algorithms used in the present and previous studies: Woolnough and colleagues obtained their measure of open nodes and closing nodes from a left-corner parsing model, which has properties of both bottom-up and top-down parsing algorithms. The present study used node counts obtained from top-down and bottom-up parsing algorithms; measures closely related to the number of parsing operations as used by Nelson and colleagues (43), a study that reported a positive relationship between open node count and HGP. While we agree with Woolnough and colleagues (44) that the choice of covariates has a nontrivial impact on uncovering neural variance related to syntactic processing, our results indicate a further dependency on the parsing algorithm used to formalize the structure of the input (18,67,68).

### 4.2. Structure building is supported by inter-regional communication

Bottom-up node counts explained variance in pars triangularis, the anterior portion of the STG and MTG, and electrodes in the dorsolateral prefrontal cortex. The positive contributions of top-down node counts were dispersed, being found in the SMG and inferior parietal cortex, the MTG, and several electrodes in pars orbitalis. This spatial distribution is in line with other findings from syntactic processing in HGP (43,44,46,47) and with earlier fMRI and lesion results of syntactic processing (74,75).

The TRFs of those electrodes revealed that HGP increases and decreases alternately between the identified regions. For bottom-up node counts, we observed recurrent fluctuations in pars triangularis and the anterior temporal lobe, with a transient increase in HGP at around 300 milliseconds post word onset, and another at around 600 milliseconds. For top-down node counts, we saw an initial increase of HGP in inferior parietal areas at around 100 milliseconds, followed by a decrease in HGP in areas further away from the sylvian fissure, and an increase in middle-to-anterior STG and MTG. Then, approximately 800 milliseconds post word onset, HGP decreases as a function of top-down node count in the electrodes in parietal cortex and around the central sulcus.

The recurrent character of these responses suggests that structure building is not an instantaneous process; rather, it supports the perspective that structure building is a process that depends upon repeated communication between two or more (distant) areas. Consistent with this proposal, gamma activity has been suggested to mediate interregional communication (37). The present finding is reminiscent of previous MEG findings of multi-stage responses to words in sentence context (76,77), and is in line with models of language comprehension that assign a crucial role to communication between the temporal lobe and other (non-adjacent) areas for structure building (2,78,79).

The patterns in the TRFs that are revealed by the clustering approach are slow in character. With two or three cycles in the time-window of 1100 milliseconds, this places them in the delta-theta range. The modulations may be related to slow neural activity. HGP has been shown to be coupled to delta phase (<4 Hz) (80) and theta phase (4-8 Hz), and theta phase-amplitude coupling could coordinate activity in distributed cortical areas (81). Hypotheses about a role of coupling between high- and low frequencies in structure building are present in several models of structure building (3,4,9). It is possible that these slow rhythms, either being shaped by the incoming acoustic input (82–84) or by the inference of linguistic representations (72,77,85,86), sensitize neural populations responsible for the transformation underlying structure building at their time-scale (80,87). This may then induce a rhythm in the neural populations that represent the computation itself (2,9). Of course, this is hypothetical, and future investigations of syntactic structure building may directly consider phase-amplitude coupling between slow oscillatory activity and HGP.

The contributions to model fit as well as the TRF clustering analysis revealed modestly spatially distinct profiles for processes associated with bottom-up and top-down node counts. While bottom-up appears to recruit two areas that are spatially close to each other and to the sylvian fissure, namely the temporal pole and IFG, top-down appears to involve synchronized decreases in areas further away from the sylvian fissure. The spatial distinction between top-down and bottom-up is reminiscent of dual-stream models of language comprehension. The spatial spread of activity associated with bottom-up node count processing maps onto a part of the ventral stream, namely the anterior temporal lobe and IFG, which are connected by the uncinate fasciculus the temporofrontal extreme capsule fasciculus (88–90). The effect of top-down node count, however, appear more in line with the sites that are connected by the dorsal stream, namely the temporo-parietal junction, the middle temporal lobe, and the IFG, which are connected via the arcuate fasciculus and the superior longitudinal fasciculus. While some models argue for a role of either the ventral stream (89,91) or the dorsal stream (92) in syntactic processing, our findings appear in line with the involvement of both pathways (93–96). The bottom-up/top-down distinction we observe here somewhat aligns with interpretations of the ventral stream subserving the building of local phrase structure, with the dorsal stream being responsible for global computations and resolving hierarchical dependencies in syntactically complex sentences (93,94). Based on these results, we would suggest that communication between cortical areas connected in a dorsal network supports predictive or top-down syntactic processing – important for the resolution of long-distance dependencies; while communication between cortical areas in the ventral network supports bottom-up syntactic processing. Due to the constraints on within-participant cortical coverage and between-participant variability in cortical loci present in datasets of ECoG recordings, complimentary evidence from (source-localized) magnetoencephalography or magnetic resonance imaging techniques is needed to evaluate this proposal.

### 4.3. A relationship between lexical probability and global disinhibition

The ‘joint encoding’ analysis revealed that surprisal was encoded in most of the recorded signals and explained the most variance, as indicated by the number of electrodes and coefficient size. The effects are largest in the posterior STG and pars triangularis and opercularis of the IFG. Entropy was encoded in the STG and IFG as well, though its coefficients were smaller than for surprisal and the positive coefficients had a more anterior distribution. The contributions of surprisal and entropy are positively correlated, suggesting that when surprisal values explain variance in the neural signal, it is more likely for entropy to do so as well. These findings suggest that lexical uncertainty and surprise tend to be represented by similar neural populations.

HGP globally increases as a function of both increasing uncertainty and surprise. The peaks are around 400-500 milliseconds in the STG/IFG, though generally the effect of entropy appears slightly later and slightly more anterior than the effect of surprisal. These effects are partially in accord with previous findings; positive coefficients for surprisal have been reported before, including the peak at 400 milliseconds (31,50), though trigram entropy has elicited negative coefficients for HGP in the 200-500ms time-window (43). This study (43) reported that syntactic features had a larger impact on model fit than the probabilistic features. This difference between studies may be at least partially attributed to the different surprisal estimates – trigram models versus GPT2-derived surprisal used here – in line with previous reports of larger contributions for probability estimates from transformer-based architectures versus n-gram-derived estimates in models of neural activity during speech comprehension (30,97).

That surprisal explained the most variance is compatible with the view that surprisal is a sum-statistic pooled over many possible sources of linguistic information (14,98), particularly when obtained from large language models, which use a large contextual window for next-word prediction. Such sources of linguistic information include syntactic representations and semantic relatedness, as well as pragmatic cues such as register, but also pure probabilistic information; i.e., how frequently linguistic units (co-)occur. This information is theoretically separable from other sources of information, as is exemplified in studies of statistical learning (22,40,99,100), but it is inherently correlated with linguistic information in speech and language. While all these sources of information can affect our response to incoming input (e.g., 101), it is to date unknown whether this change in response hinges on probability. In other words, because of the nature of probabilistic features, it is unclear whether all effects associated with them are reflections of probabilistic processing or representation of probability. Given the evidence for statistical learning and the use of statistical cues in situations such as ambiguity (102), we assume that at least some part of the variance captured with surprisal and entropy is probabilistic processing in its purest sense; the rest of the variance may reflect constraint from other available cues (31). The current study cannot disentangle those sources of variance.

We interpret the time courses of the effects of surprisal and entropy on HGP as follows. The state of neural activation at any time point is affected by previous input, constituting a structuring context for the processing of incoming stimuli. Here, we propose that the effects observed are reflections of global, transient changes in the neural state that are provoked by the processing of new input, inhibiting some neural populations more than others to minimize competition between related- and previously inferred representations (2). The processing of new input changes this state. Being sum-statistics over many potential linguistic representations, surprisal and entropy are predictors that are apt to capture such global changes. In contrast to low-surprisal words, high-surprisal words are unexpected given previous input, and neural populations coding for these words were inhibited; more excitatory (or disinhibitory) activity will be necessary to reach word recognition. Consequentially, the recognition of unexpected words will produce a larger change in the sensitization pattern of neural populations. For entropy, a similar mechanism might be at play. Recognizing words in positions of high uncertainty about the continuation – i.e., a relatively unconstrained path – will lead to a larger change in the current state of the brain than recognizing words in constrained condition, provided that the word matches the expectations (low surprisal).

### 4.4. Speech comprehension is a flexible weighting of cues

Besides the encoding of individual features, we also set out to examine how they were jointly encoded, and whether they affect each other. To do so, we evaluated whether the contributions to model fit were correlated between features, and whether the neural encoding of syntactic structure is affected by the probability of a word. Correlations between feature coefficients from the joint encoding analysis indicated that the contribution from bottom-up node counts is smaller when entropy and surprisal explain more variance. Bottom-up node count coefficients correlate negatively with both probabilistic features. This suggests that bottom-up structural processing draws more on neural populations that do not strongly encode contextual lexical probability; a finding that is in line with previous studies reporting spatial distinctions between lexical and syntactic conditions (46,47). At the same time, this analysis revealed that top-down syntactic information is more likely to have a greater impact when entropy does, too. This suggests that structural predictive information and lexical uncertainty are more likely to be represented by the same (or minimally *very* closely situated) neural populations. While this could simply indicate a joint representation, their co-occurrence could also signal a process in which (A) lexical uncertainty determines the weighting of predictive structural information or (B) predictive structural information determines the relative importance of lexical uncertainty.

We find interactions between probability and syntax in several electrodes. In the electrodes that display an interaction between entropy and top-down, the time-course of HGP modulation by top-down node counts appeared to be shaped by low-entropy words more than by high-entropy words. This suggests that lexical uncertainty affects the encoding of top-down structural information. Considering also the positive correlation between the coefficients of these features, we suggest that lexical uncertainty modulates the weighting of anticipatory structural information such that it plays a larger role in shaping the comprehension process when lexical uncertainty is low; a trade-off in which predictive cues receive stronger weighting in the comprehension process in low-uncertainty states. Evidence for a trade-off at this level adds to previous work that has shown that high word entropy leads to increased low-frequency tracking of lower-level features, such as acoustic and phonemic features (103,104).

In the electrodes in which bottom-up structure building interacts with surprisal, the TRFs revealed that these neural populations are more sensitive to bottom-up structure building when surprisal is high. This suggests that these neural populations enhance structural representations in states of low word probability. Both surprisal and bottom-up node counts are ‘reactive’ features, in the sense that they reflect information gain after the word has been observed. Finding a larger effect for a syntactic feature of the reactive type for high surprisal words is in line with either a lack of anticipatory syntactic information (high entropy, high surprisal), or a lack of *correct* anticipatory syntactic information (low entropy, high surprisal), both of which require reliance on bottom-up structure building to integrate the word into the sentence. These interpretations are in line with dynamic accounts of language comprehension in which cues are weighted flexibly (105–111).

The short duration of the recordings used in this study decreases the reliability of the TRFs estimated in the split-feature approach and, by consequence, increases the likelihood of null results. Despite the low number of electrodes identified here, these results provide compelling, if preliminary, evidence that the process of structure building as captured by HGP is mediated by lexical uncertainty and surprise. Future research on longer recordings should provide a more detailed view of the direction of the interaction between top-down and entropy and aim to elucidate the direction of the interaction between bottom-up structure building and lexical surprisal; the effects may very well be bi-directional.

## 5. Conclusion

This study provides evidence for an interactive, dynamic process of speech comprehension that continuously integrates syntactic and lexical probabilistic information. The results show that local, high-frequency (70-150 Hz) cortical activity jointly represents lexical probabilistic information and syntactic structure, and that these features are actively integrated during spoken language comprehension. The current results indicate that high gamma power is sensitive to multi-word estimates of constituency structure and suggest that syntactic structure building depends on communication between interconnected regions of the dorsal- and ventral streams of speech processing. Furthermore, we provide evidence that bottom-up syntactic information is less likely to be encoded by neural populations that strongly code for lexical probability measures, while top-down syntactic information shares neural resources with lexical uncertainty. We propose that lexical uncertainty modulates the weighting of anticipatory structural information, with these cues being interactively resolved in a distributed and often overlapping network of perisylvian cortical patches.

