## Supplementary figures for "Linguistic structure and probability are jointly encoded in high gamma power"

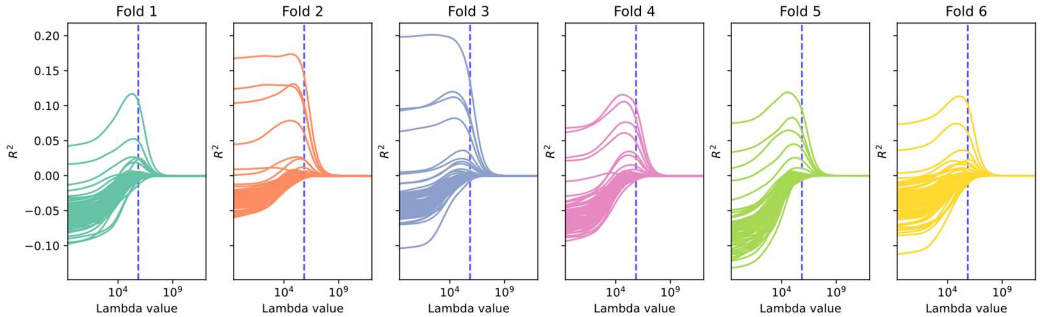


Figure S1. Reconstruction accuracy ($R^2$) values for various lambda values across folds for one participant (sub-05) for the model including all features. Each panel reflects a fold. In each panel, the individual lines reflect one electrode. The vertical, blue dashed lines indicate the selected regularization parameter.

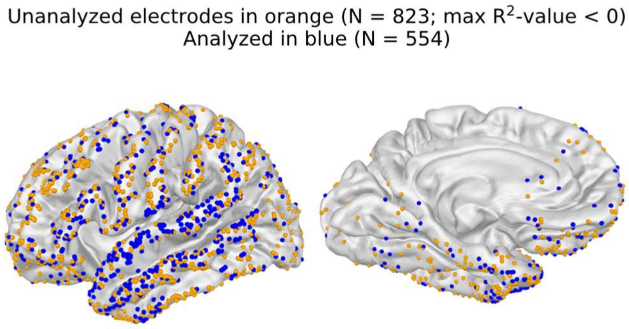


Figure S2. Unanalyzed electrodes (orange) and analyzed electrodes (blue). The electrodes that were skipped in analysis were those that had a negative value as its largest $R^2$-value (out of all the fitted main-effects models).

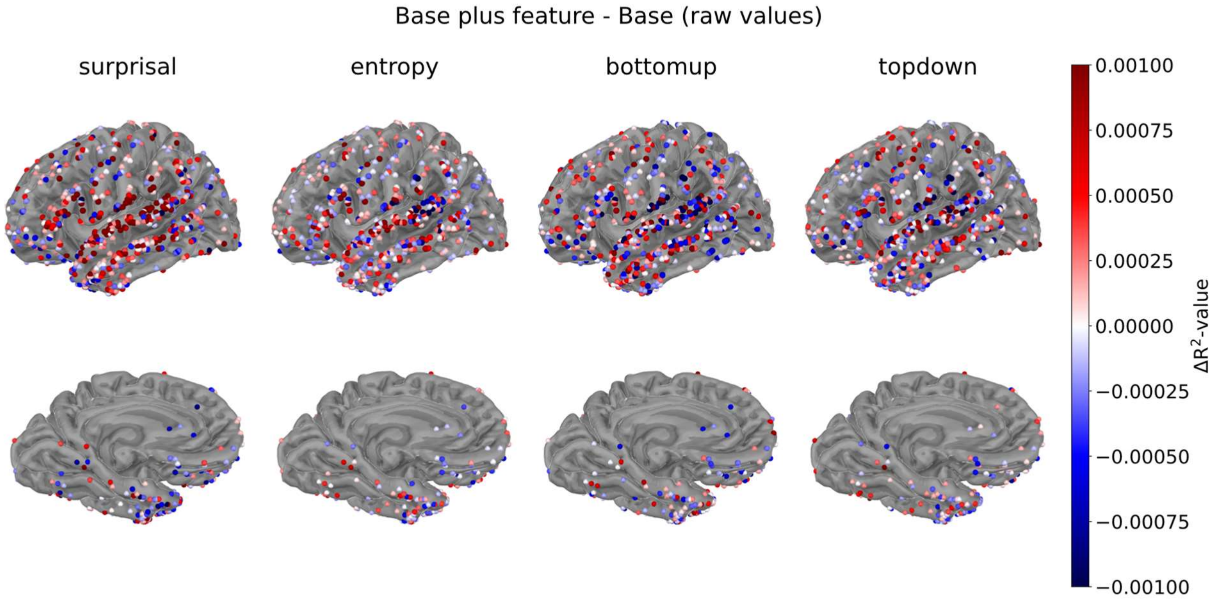


Figure S3. Contrast between raw $R^2$-values from the base model and the model that contained the feature under scrutiny as its only extra feature (base + 1). For each feature, the base model $R^2$-values were subtracted from the model with the feature. Positive effects mean that the model with the feature was better than the model without the feature.

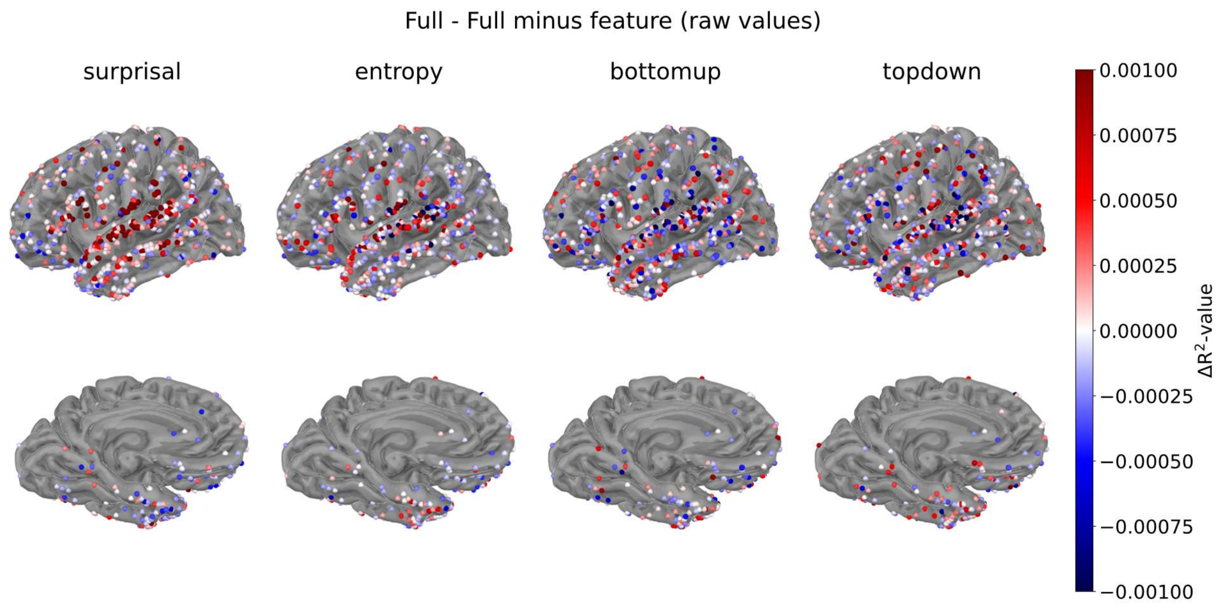


Figure S4. Contrast between raw $R^2$-values from the full model and the model that did not contain the feature under scrutiny as the only missing feature (full - 1). For each feature, the $R^2$-values of the model that did not have the feature were subtracted from the full model. Positive effects mean that the model with the feature was better than the model without the feature.

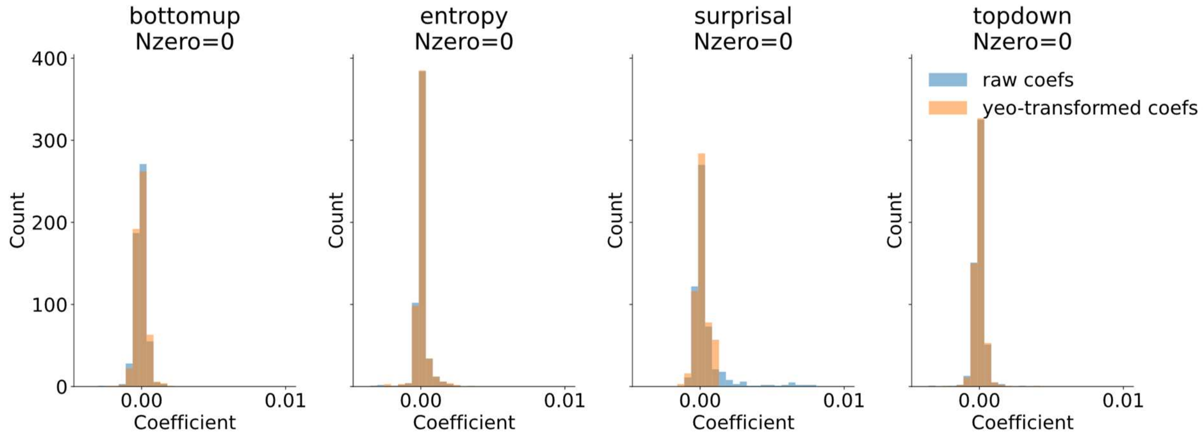


Figure S5. Untransformed and Yeo-Johnson transformed coefficients ('joint encoding' analysis) for each feature.

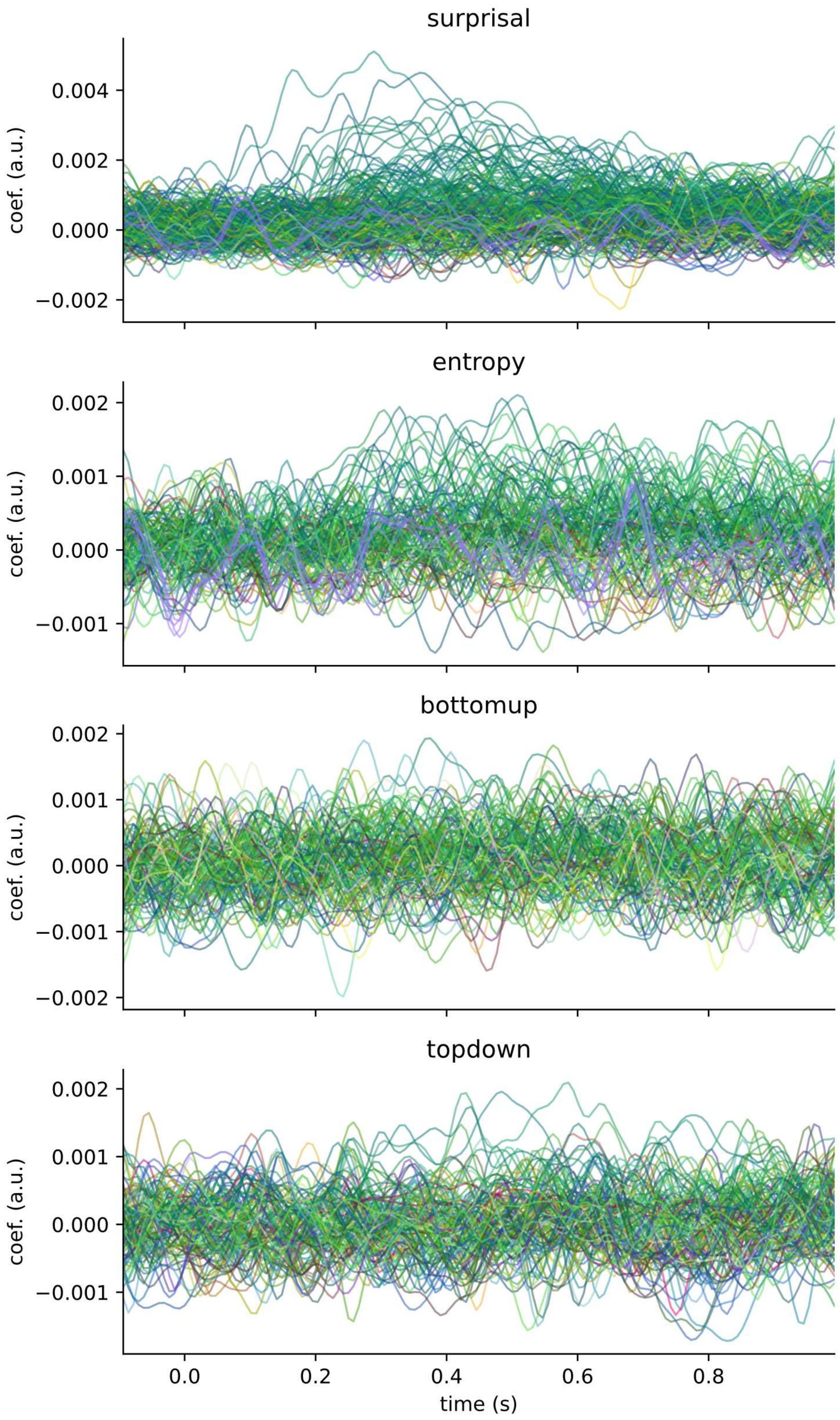


Figure S6. Unsmoothed TRF time courses for each feature. RGB colors map onto the same topography as accompanying Figure 11 in the main text.

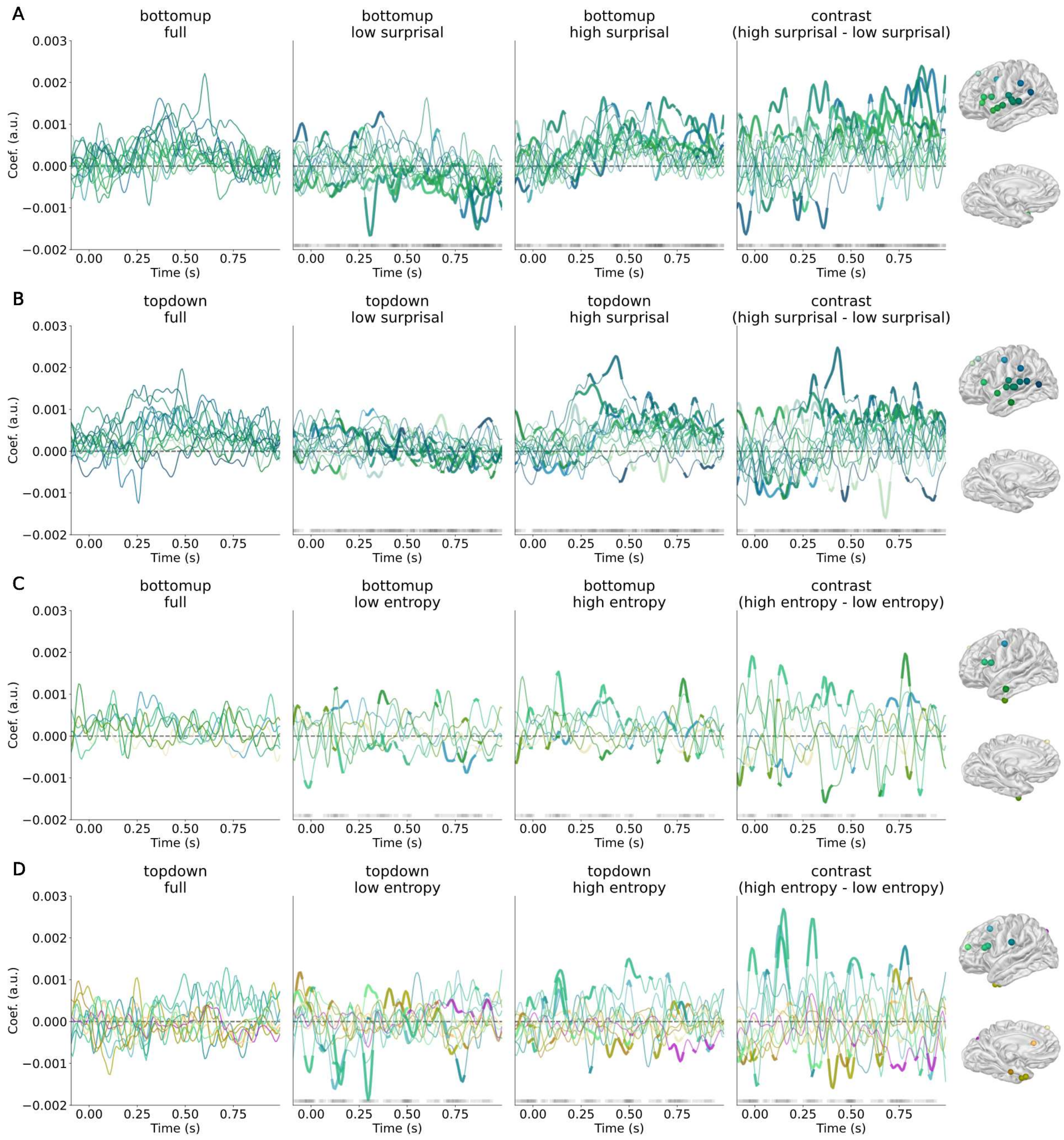


Figure S7. Unsmoothed Temporal Response Functions for the split features; accompanies Figure 13 in the main text. (A) Bottom-up split by surprisal; (B) Top-down split by surprisal; (C) bottom-up split by entropy; (D) top-down split by entropy. Only time courses for significant electrodes are displayed. The first column shows the TRF for the feature trained on all the data (no split) for the same sensors. The middle two columns show the halves of the split feature. The last column displays the contrast. Sensors and time-points in bold were those that were in the upper- or lower 2.5% of the distribution of the difference between 1000 random splits. The grey bar at the bottom shows the time-points; darker grey means more sensors were significant at that time-point. RGB colors indicate sensor position.
